# Individual Variability in the Innate Functional Organization of the Human Brain

**DOI:** 10.1101/2021.03.24.436788

**Authors:** M. Fiona Molloy, Zeynep M. Saygin

## Abstract

The adult brain is organized into distinct functional networks, forming the basis of information processing and determining individual differences in behavior. Is this network organization genetically determined and present at birth? And what is the individual variability in this organization in neonates? Here, we use unsupervised learning to uncover intrinsic functional brain organization using resting-state connectivity from a large cohort of neonates (Developing Human Connectome Project). We identified a set of symmetric, hierarchical, and replicable networks: sensorimotor, visual, default mode, ventral attention, and high-level vision. We quantified individual variability across neonates, and found the most individual variability in the ventral attention networks. Crucially, the variability of these networks were not driven by SNR differences or differences from adult networks (Yeo et al., 2011). Finally, differential gene expression provided a potential explanation for the emergence of these distinct networks and identified potential genes of interest for future developmental and individual variability research. Overall, we found neonatal connectomes (even at the voxel-level) can reveal broad individual-specific information processing units. The presence of individual differences in neonates and the framework for personalized parcellations demonstrated here has the potential to improve prediction of behavior and future outcomes from neonatal and infant brain data.

## Individual Variability in the Innate Functional Organization of the Human Brain

Elucidating the organization of the brain has been a major objective in neuroscience for many decades. By understanding the arealization of the brain, and how brain areas connect and communicate with one another, neuroscientists hope to better understand information processing and how the brain performs complex mental functions. Early approaches to understanding the human brain relied on postmortem samples and similarities in cell types and myelination were used to distinguish between brain areas. Brodmann’s areas (Brodmann, 1909), identified via cytoarchitecture, are still widely used to label and identify brain regions in neuroimaging studies. Advancements in noninvasive neuroimaging brought about newer atlases which are based on differences in structure and tissue composition that can be identified with MRI. As opposed to early approaches which were based on postmortem samples from one or a few individuals, these atlases can be identified in vivo and are typically based on structural information from a large group of individuals, thus increasing the applicability of these atlases to the general population.

More recently, resting state fMRI scans (measuring spontaneous brain activity) show that the brain is organized into functional networks that support distinct perceptual and cognitive functions e.g., visual, default, limbic, somatomotor, ventral attention, dorsal attention, and control networks (Yeo et al., 2011). These networks were identified by measuring co-fluctuations of BOLD signal during rest (i.e., functional connectivity to the rest of the brain) and clustering brain regions together based on the similarity of this functional connectivity profile.

Numerous studies support these conclusions, identifying similar network structures via functional connectivity in the adult cortex (Choi et al., 2018; Power et al., 2011; Smith et al., 2009) and also in the cerebellum (Buckner et al., 2011). There is evidence that this network organization is heritable (Ge et al., 2017; Glahn et al., 2010) and reveals important individual differences that are predictive of individual variability in behavior for both typical individuals (Kong et al., 2019; Rosenberg et al., 2016) and for mental illness (Buckholtz & Meyer-Lindenberg, 2012; van den Heuvel & Sporns, 2019). The importance of individual variability in both brain organization and behavior has demonstrated the need for individual-level parcellations (Bijsterbosch et al., 2020). In adults, there exist numerous individualized approaches of varying resolutions based on functional connectivity (Blumensath et al., 2013; Hacker et al., 2013; Kong et al., 2019), surface-based anatomy (Desikan et al., 2006), or multiple modalities (Glasser et al., 2016); (see Eickhoff et al., (2018) & Arslan et al., (2018) for comprehensive reviews and comparisons for adults). But the presence of a structured, heritable functional organization and its link to individual variability in behavior in adults brings us back to a fundamental question in neuroscience and psychology: is the organization of the human brain innate and what individual variability in this organization exists at birth?

The human brain undergoes major changes over the first few years of life, including large overall growth and expansion due to e.g. synaptogenesis and myelination, with certain brain regions developing earlier than others (Gogtay et al., 2004; Kinney et al., 1988; Lebel et al., 2008). And yet humans are born with a rich set of mental faculties, presumably supported by existing neural architecture that is genetically determined prenatally (Dehaene-Lambertz & Spelke, 2015; Price et al., 2006; Takahashi et al., 2012). Infant neuroimaging techniques have flourished over the past few years and there is now evidence that young infants display neural signatures for phonetic representations (Dehaene-Lambertz & Gliga, 2004) and sophisticated visual functions such as face and scene processing identified with fMRI and fNIRs (Deen et al., 2017; Farroni et al., 2013); moreover, recent work shows the distinct connectivity patterns present at birth may determine the location of this high-level visual cortex (Kamps et al., 2020; Li et al., 2020; Saygin et al., 2016). Previous work using resting-state connectivity during natural sleep in preterm infants suggests large-scale organization is present prenatally by the 3^rd^ trimester (Doria et al., 2010); this and other work in infants within the first year of life have identified networks that closely resemble those found in adults using independent components analyses (Doria et al., 2010; Fransson et al., 2007, 2011; Gao et al., 2009; Liu et al., 2008; Smyser et al., 2011) and graph theoretical approaches (Ball et al., 2014; Shi et al., 2018).

However, these methods required a priori of interest or user-specified components/ networks to define the connectome and/or to constrain the boundaries of networks. We apply no constraints based on location or known adult networks, which allows us to uncover both localized and distributed neonate-specific networks based on neonatal resting state connectivity alone. Additionally, instead of excluding components based on lower intersubject agreement, we quantify within-subject agreement of these networks and identify and quantify networks with high variability across subjects. Crucially, while individual differences have been found to alter underlying networks in adults (Kong et al., 2019), it is unclear if, and to what degree, individual variability in brain networks exists at birth. Consequently, we describe a framework to identify individual networks, in addition to the group-level parcellations. To the best of our knowledge, a fine-grained (i.e., voxel by voxel), whole-brain approach to cluster resting state data in a large group of newborns has not yet been attempted. Can we use voxelwise connectivity to delineate the intrinsic network structure of the neonate brain? Are these networks distributed and symmetrical like they are in adults? Are they variable among individuals, and what are the underlying differences in genetic expression that may determine this organization prenatally?

We aim to answer these remaining questions about the functional organization of the brain at birth using a data-driven, high resolution unsupervised learning approach (see **Figure 1** for an overview). We use the voxel-wise resting state data from the developing Human Connectome Project (dHCP; Hughes et al., 2017); https://www.developingconnectome.org) on 267 term-birth newborns and determine voxel-to-voxel functional connectivity. Following a similar approach as (Yeo et al., 2011), these data are then clustered based on k-means unsupervised clustering. We compare training and independent test set clustering solutions for k = 2 to k = 25 networks and find the optimal solutions on the whole brain (subcortex and cortex) and separately for the cortex only (for direct comparison to the adult clustering solutions from (Yeo et al., 2011). We then evaluate individual variability of the resulting networks on the personalized parcellations in the independent test set. Finally, we explore the possible genetic basis of these networks by matching our networks to the genetic data available from the Allen Human Brain Atlas (AHBA; Hawrylycz et al., 2012).

**Figure 1:**
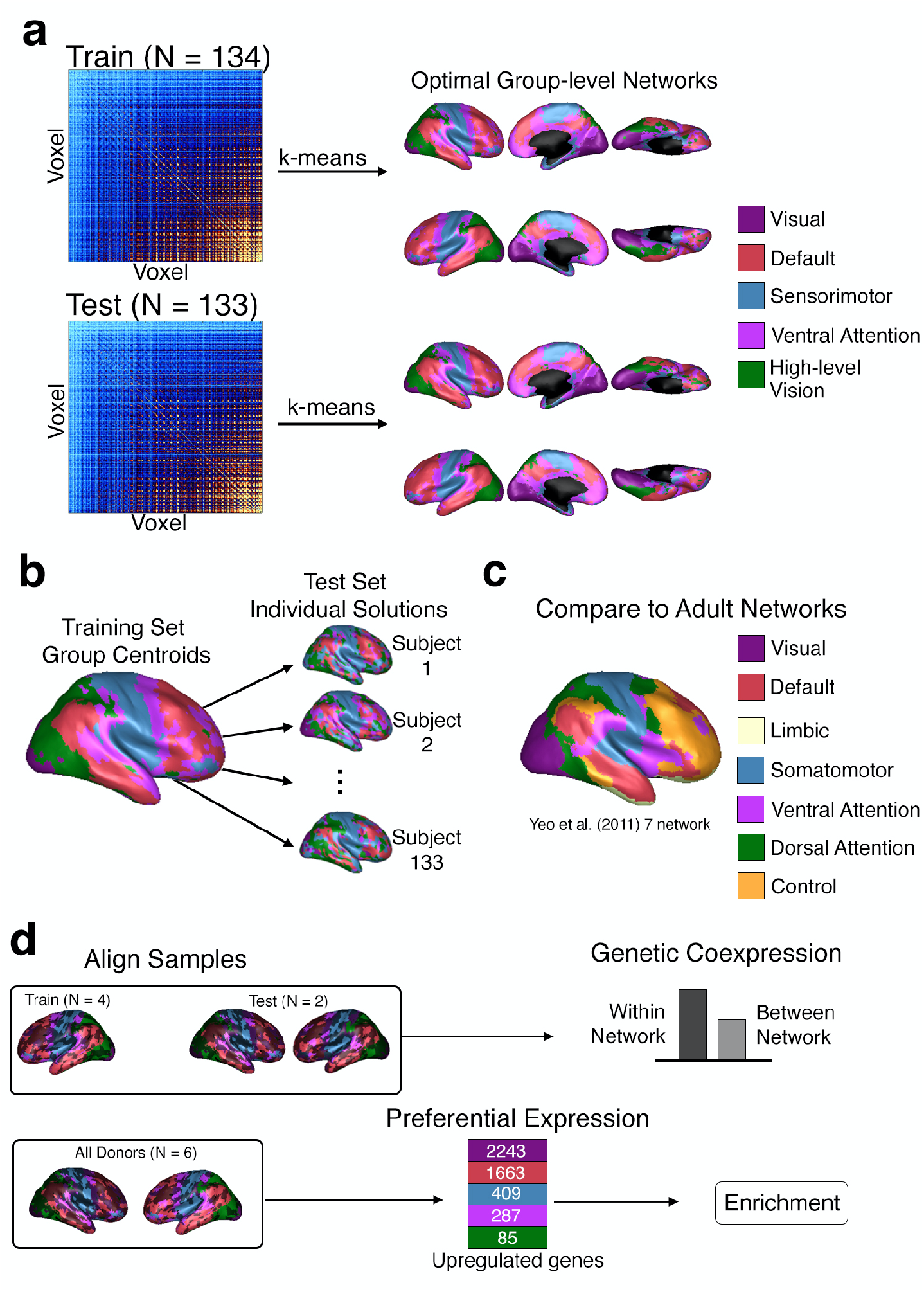
Overview of Analyses. **a.** We split neonate data into two randomly defined groups (train and test sets) and calculated voxel-to-voxel functional connectivity (illustrated by the example connectivity matrices). We then calculated clustering solutions (from k = 2 to k = 25) on the voxel-by-voxel neonate connectome for the training and test sets independently. We identified the optimal networks (optimal k- clusters) based on the best overlap of the train and test solutions. **b.** To investigate individual variability of these networks, we generated individual solutions for each individual in the test dataset using the group-level centroids from the training group solutions (5-network solution pictured, with three individual subject results). **c.** We compared group-level neonate networks to the group-level adult networks from Yeo et al. (2011) and quantified overlap. **d**. Genetic expression of the networks was explored by aligning samples from the Allen Human Brain Atlas (AHBA) to the 5-network neonate solution. We first split the AHBA samples into train and test groups to explore within and between network genetic expression. We then combined all available data across AHBA donors to find preferentially expressed genes by network.

## Results

### Optimal Solutions

First, we determined neonatal networks underlying high-resolution resting-state connectivity data (**Figure 1a**). We calculated voxel-to-voxel connectivity for the training set (134 neonates) and test set (133 neonates) of neonatal resting state data (mean age at scan: 1.31 weeks). These connectomes were clustered based on similarities of voxel-wise connectivity profiles, starting from k = 2 to k = 25. We then chose the optimal number of networks/ clusters, k, by identifying the solutions that best fit the neonatal data (quantified by the silhouette index, SI) and were reliably reproduced (quantified by the overlap between the independently calculated test and train solutions). The clustering analysis revealed two optimal solutions: k = 5 and k = 8 (**Figure 2a**). These k-solutions produced parcellations that maximized within-network similarity, while minimizing between-network similarity, i.e., these solutions had high SIs (mean: 0.81 for k = 5 and 0.90 for k = 8). These networks were replicated in the independent test group, indicated by the high overlap between the test and train solutions (mean dice coefficient: 0.89 for k = 5 and 0.88 for k = 8; Adjusted Rand Index: 0.73 for k = 5 and 0.72 for k = 8).

**Figure 2.**
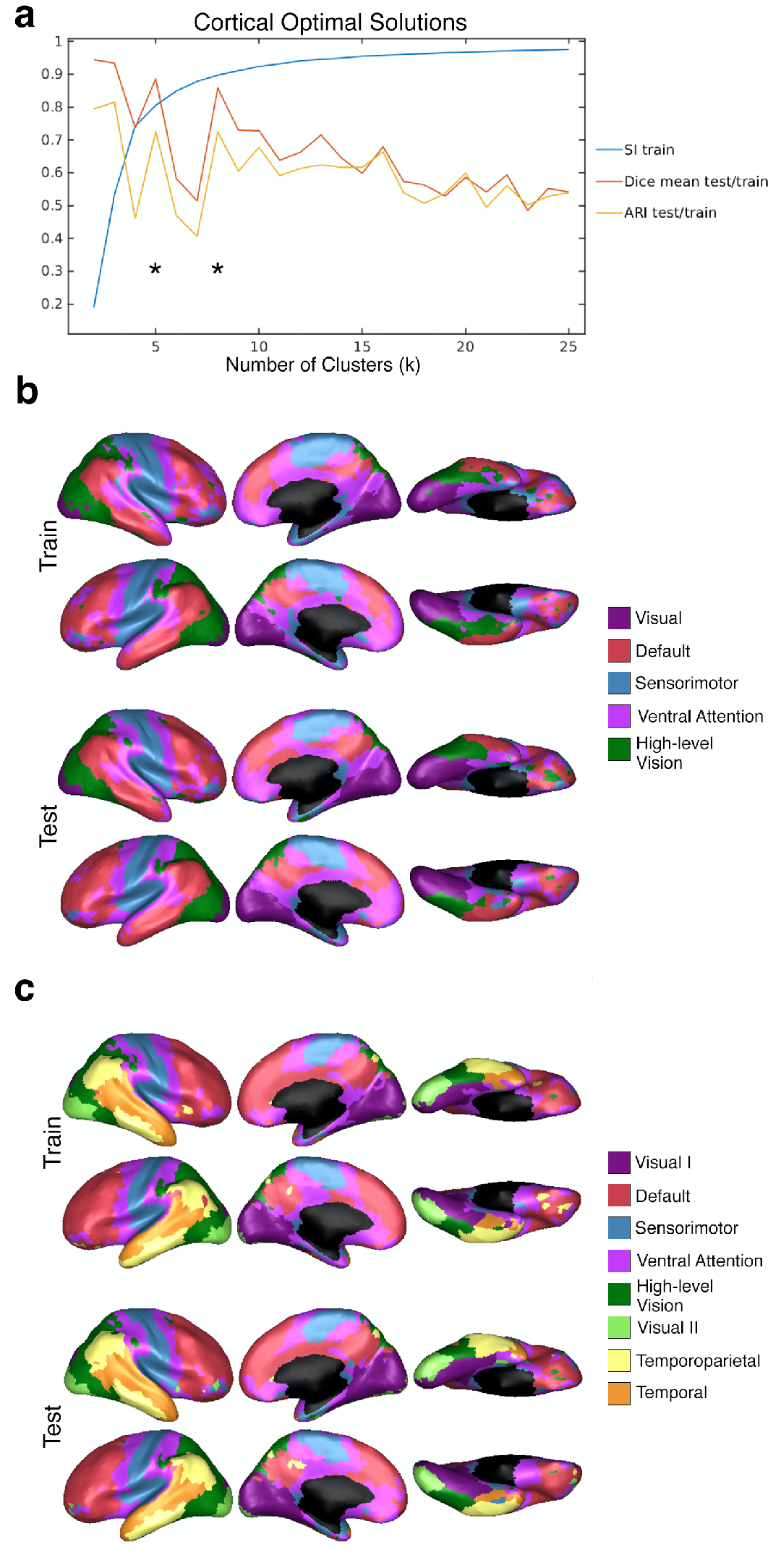
Optimal Cortical Solutions. **a**. Different quantitative evaluations (y-axis) of clustering solutions for each of the k-clusters (x-axis). The blue line shows the Silhouette Index (SI) for clustering in the training set (essentially the fit of the clustering solutions to the underlying connectivity data). The red and yellow lines show the overlap of the training and test set solutions quantified by the Dice coefficient (red line) and Adjusted Rand Index (ARI; yellow line). **b**. Optimal k = 5 solution for the training and test sets. **c**. Optimal k = 8 solution for the training and test sets. b and c illustrate the hemispheric symmetry of the solutions and similarity of the solutions between train and test sets.

The 5-network optimal solutions for the training and test sets are shown in **Figure 2b**. Despite the lack of structural or spatial constraints, network components were spatially continuous and symmetric. Approximately 80% of the voxels were symmetric (i.e., had the same network assignment in the corresponding voxel across hemispheres): 80.31% for the training solution and 80.63% for the test solution. The networks corresponding to the visual and sensorimotor networks (purple, green, and blue) were quite spatially localized. Interestingly, these networks were the first to be differentiated at lower k values and were further subdivided as k increased. At k = 2, the visual and sensorimotor networks formed one network separated from the rest of the cortex. With k = 3, the visual and sensorimotor networks separated into distinct networks. With k=4 and 5 (optimal solution), the visual network split into low-level visual and high-level visual clusters; higher k solutions (including k=8, the other optimal solution, discussed below) revealed a further split of the visual network into anterior/posterior components. The hierarchical emergence of networks is described in more detail (**Supplementary Materials**) and illustrated in **Supplementary Figures 11—12 and Supplementary Movie 1**.

For the optimal cluster solution of k=5, the blue cluster, which we refer to as the sensorimotor network, covers the somatosensory cortex, the motor cortex, and parts of the medial temporal lobe. Two of the clusters span the visual cortex. The purple cluster includes much of the occipital cortex and is therefore named the (early) visual network. The other visual network (green cluster in **Figure 2b**) strikingly seems to encompass high-level visual cortex spanning the dorsal and ventral visual streams (including occipital and parietal/temporal cortices) and is therefore referred to as the high-level vision network. The remaining clusters (red and magenta) were more distributed across the brain and were named based on qualitative similarities to adult networks (Yeo et al., 2011). The default network (red cluster in **Figure 2b**) contains frontal, superior temporal, and superior/parietal regions. Finally, the ventral attention network (magenta cluster in **Figure 2b**) is comprised of the frontal eye fields and cingulate gyrus.

The higher-resolution optimal clustering solution revealed eight distinct networks (**Figure 2c**). In line with the aforementioned hierarchical emergence of additional networks, the network borders remained roughly the same as that observed in the 5-network solution, but additional clusters emerged in occipitotemporal and parietal regions. In the same manner as the lower resolution solution, the network components were continuous and symmetric (78.76% matching voxels for the training solution and 80.76% matching voxels for the test solution). The ventral attention and sensorimotor networks were highly preserved between the 5-network and 8- network solutions. The other networks in the 8-network solution further segmented the temporal and occipital lobes. The low-level vision network in the 5-network solution was further split into a medial network (purple, Visual I) and a lateral network (light green, Visual II). An additional temporoparietal network (yellow) was distinguished from the lateral posterior components of the default network. Finally, a temporal network (orange) emerged in a region that consisted of the ventral attention and default network in the 5-network solution.

### Individual Differences

How are these optimal networks affected by individual differences in connectivity? To answer this, we found individual solutions for the 133 neonates in the testing set, assigning a network for each voxel (for each individual) based on how well that voxel’s connectivity profile fit the group-level 5-network solution of the training group (**Figure 1b, see individual examples in Figure 3a**).

**Figure 3.**
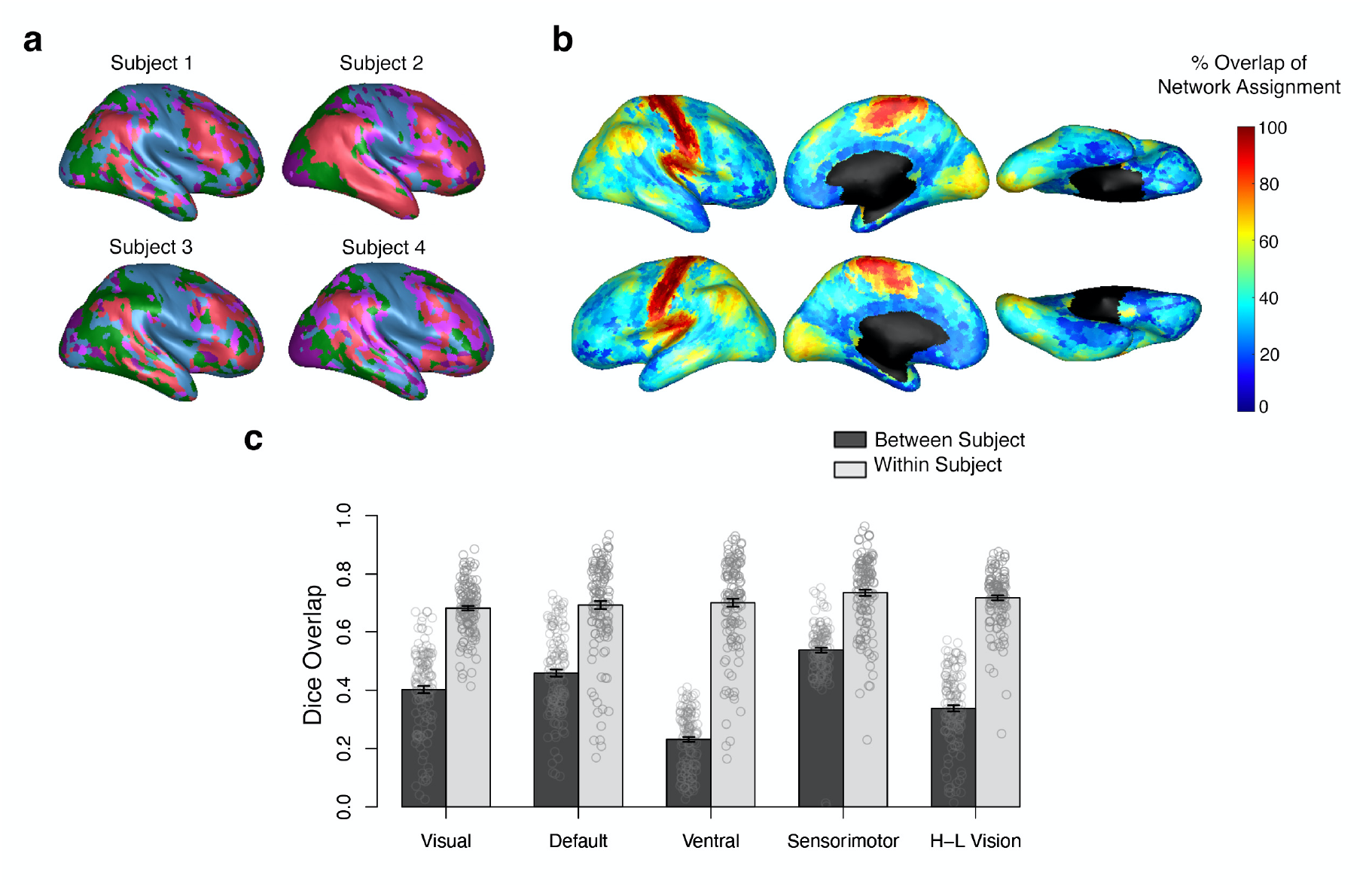
Individual Differences in the 5-network Solutions. **a.** Example individual solutions for four randomly selected neonates. **b**. Voxel-wise between-subject proportion of overlap for the 5 networks. Voxels in warmer colors depict a higher proportion of individuals who were assigned to the same network. **c.** Within-subject overlap (mean: dark bars, individuals: open circles) and between-subject overlap (mean: light bars, individuals: open circles) for all voxels within a network. For all networks, the within subject overlap is significantly higher than the between subject overlap (paired Wilcoxon signed-rank p ≤ 0.001 for all networks). The error bars indicate ± 1 standard error of the mean.

The between-subject agreement was defined as the overlap of network assignment averaged across all 133 subjects for each voxel (**Figure 3b, all optimal solutions in Supplementary Figure 14**). The between-subject overlap varied among voxels (mean = 43%, standard deviation = 20%), showing some voxels with very high between-subject overlap (100% of individuals with the same network classification) and voxels with very low between-subject overlap (0.75% of individuals with the same network classification). We found the highest between-subject overlap in the sensorimotor network, which coincides with similar findings in adults (Kong et al., 2019). Moderate between-subject agreement was found in the medial components of the visual network, and in the parietal component of the default network. The network with the lowest between-subject overlap was the ventral attention network. These voxel-wise results were corroborated using the dice coefficient per network (**Figure 3c**) showing a similar order of individual variability as the voxel-wise results, with the highest between-subject overlap for sensorimotor areas (mean dice = 0.54 ± 0.0085), and lowest for the ventral attention network (mean dice = 0.23 ± 0.0086). To ensure low overlap was not a result of noise or signal-to-noise ratio differences across the brain, we calculated within-subject overlap of network solutions by splitting up each neonate’s connectivity data into three 10-minute resting state timeseries and generating three independent connectome solutions per subject. **Figure 3b** shows the mean overlap within each network for between-subject overlap, and within-subject overlap. Across all networks, within-subject similarity was higher than the between-subject similarity (non-parametric paired two-sided Wilcoxon signed rank p ≤ 0.001 for each network, Cohen’s d > 0.80 for each network, **Supplementary Table 2**). Moreover, individual variability found in the ventral attention network does not appear to be driven by higher noise in within that network, as the within-subject variability does not differ from other networks (two-sided Wilcoxon signed rank V = 4819, p = 0.41, d = -0.012, 95% CI [-0.019,0.042] for within-subject overlap in ventral attention versus all others, compared to V=41, p = 3.7×10^-23^, d = -1.2, 95% CI [-0.22,- 0.18] for between-subject overlap in ventral attention versus all others).

### Comparison to Adults

We next directly compared the group-level optimal network solutions between neonates and those previously identified in adults. **Figure 4a** shows the 5-network neonate solution found here and the adult 7-network solution (Yeo et al., 2011). Many of the networks identified in neonates resemble those in the adults including (as labeled in the Yeo atlas): visual, default, somatomotor, ventral attention, and dorsal attention networks. However, there were no neonate networks matching the control or limbic adult networks (see **Supplementary Figure 15** for a quantitative comparison). This suggests that regions in the limbic and control networks lack discrete connectivity patterns in neonates such that they cannot be easily separated from the rest of the cortex.

**Figure 4.**
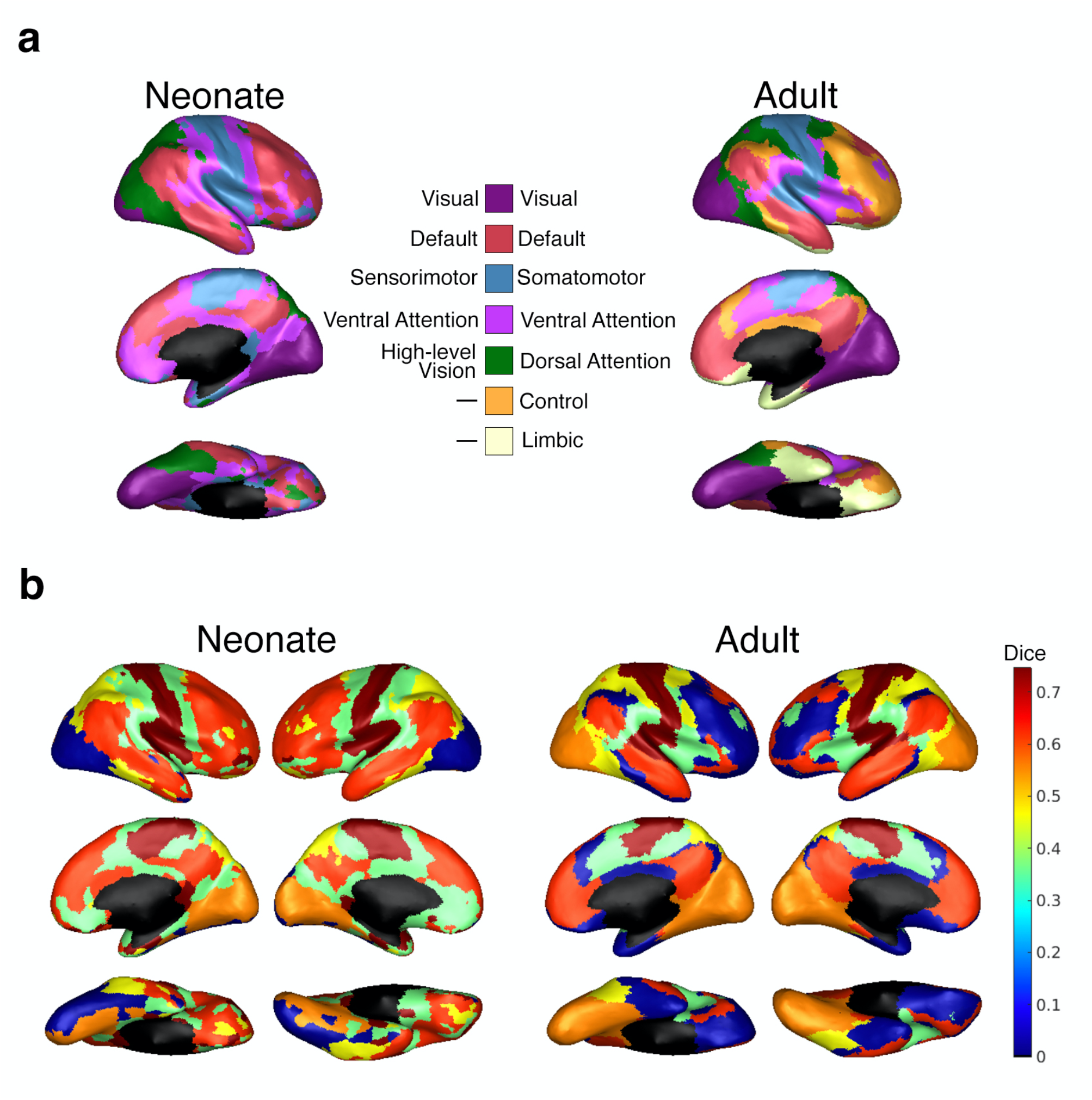
Comparison to Adult Networks. **a.** Lowest optimal solution for neonates (k = 5 networks) and for adults (k = 7 networks, from Yeo et al., 2011). **b**. Dice overlap between matched pairs of neonate (k = 7) and adult (k = 7) networks. Warmer colors show that a network is more similar between neonates and adults. The results are plotted onto the neonate networks (left) and the adult networks (right).

However, it is plausible that these networks could emerge if the total number of networks in neonates is assumed to be the same as the total number of networks in adults. To test this, we next compared the 7-network solution for neonates to the 7-network solution in adults (**Figure 4c**). The highest overlapping networks were the sensorimotor network (dice = 0.75) and the default network (dice = 0.62). Interestingly, the dice coefficients for the adult control and limbic networks remain very close to zero (control = 0.000302 and limbic = 0.00985). Thus, even at higher k-solutions, there were no neonatal networks that resembled the adult control and limbic networks. Instead, the two vision-related networks in the neonate 5-network solution (high-level vision and vision) were further split, yielding four visual networks in the 7-network solution for neonates (see also **Supplementary Figure 12** for illustration of hierarchy of networks with increasing k).

### Genetic Expression

The results so far suggest that prenatal mechanisms determine cortical connectivity such that these distinct functional networks are already present at birth, even in the absence of rich postnatal sensory experiences. What are the potential genetic differences driving these networks? Previous work revealed distinct genetic expression profiles for cortical networks in adults. We followed similar analyses using tissue samples from 6 donors available through the Allan Human Brain Atlas (AHBA) to explore the correspondence of the neonatal networks and genetic expression (**Figure 1d**). The first analysis determines how well gene sets preferentially expressed in one network account for genetic variability within that network as compared to others. The second analysis relates the network-specific gene sets to e.g. putative biological and cellular functions via gene list enrichment.

In the first analysis, to characterize differential expression by network, we assigned samples to each network from: i. four donors to define network specific gene sets and ii. two donors to independently test how well those gene sets account for within and between network genetic expression (**Figure 5a**). Preferentially expressed gene sets for each network included genes with a significant (Bonferroni-correction q ≤ 0.01) positive log2 fold-change expression in that network versus the average of all others. We used the samples in the independent donors (remaining two donors) to compare within- and between-network expression of these gene sets (**Figure 5b**). Overall, within-network expression (mean = 0.33±0.023) was higher than between-network expression (mean = 0.12±0.011; F(1,1873)=123.99, p < 2.2×10^-16^, η_p_^2^=0.06, 95% CI [- 0.24,-0.17]). There was a significant difference within- vs. between-network expression for genes in the visual (F(1,373.28)=102.92, p = 2.2×10^-16^, η_p_^2^=0.22, 99% CI [-0.58,-0.35]), default (F(1,373.11)=34.18,p = 1.1×10^-8^, η_p_^2^=0.08, 99% CI [-0.27,-0.10]), ventral (F(1,372.62)=13.41, p = 2.9 ×10^-4^, η_p_^2^=0.03, 99% CI [-0.30,-0.052]), high-level vision networks (F(1,373.26) = 32.81, p = 2.1×10^-8^, η_p_^2^=0.08, 99% CI [-0.47,-0.18]), and a trend for the sensorimotor network (F(1,3.518)=5.24, p = 0.093, η_p_^2^=0.60, 99% CI [-0.30,0.11]).

**Figure 5.**
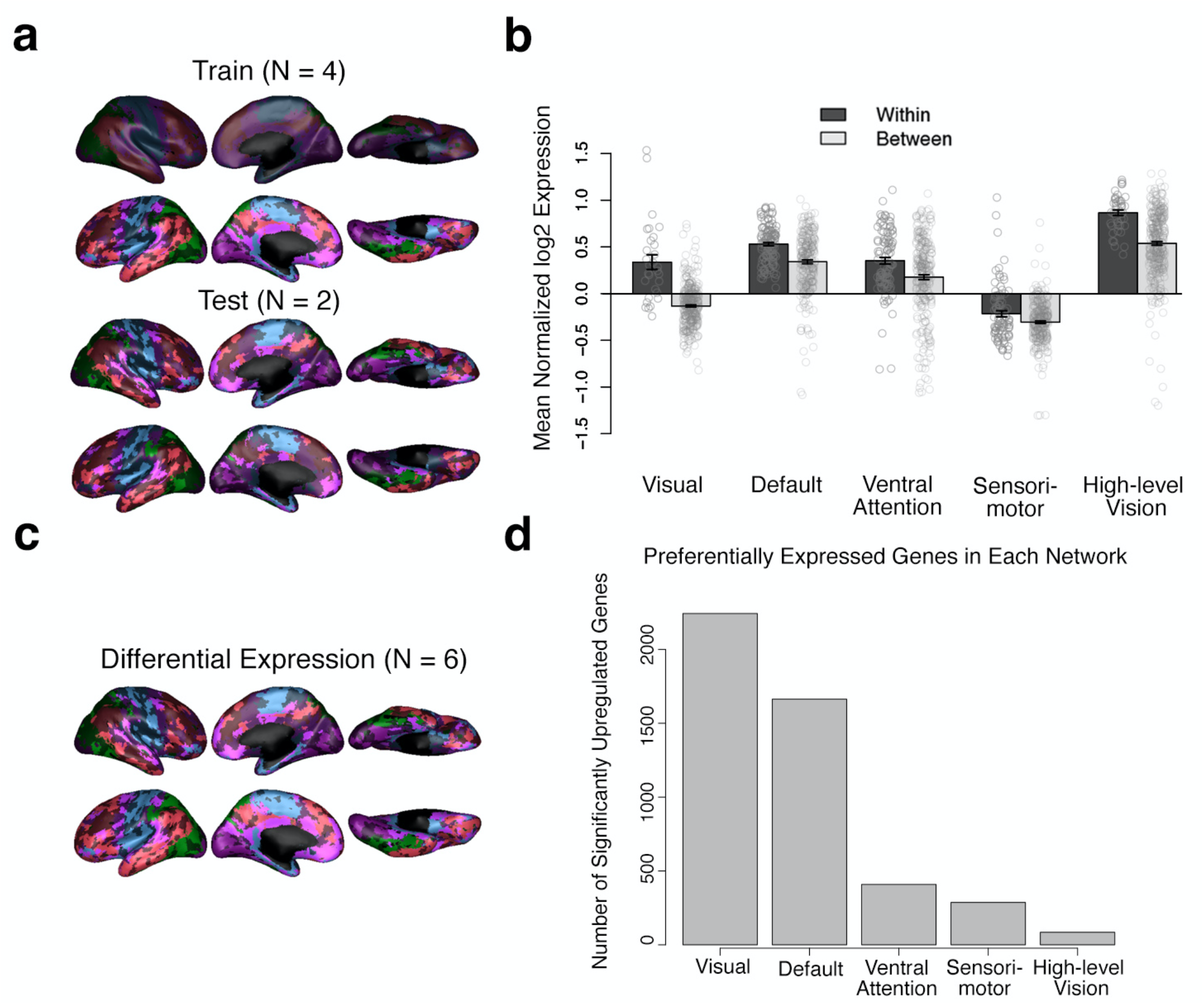
Genetic Expression by Network. **a.** Location of samples matched from the 4 left-hemispheric donors (top: used to define the network-specific gene sets) and the 2 bihemispheric donors (bottom: used to test the genetic expression within and between each network). Black-shaded regions indicate there was no matching sample/ no available donor data for that region. **b.** Genetic expression of each network’s preferential genes (identified from the 4 training donors) for within-network samples (mean: dark bars, observations: open circles) and between-network samples (mean: light bars, observations: open circles) for the 2 test donors. The error bars show ± 1 standard error of the mean. **c.** Location of samples matched to data from all available donors. **d.** Number of significant preferentially expressed genes (Bonferroni corrected p-value < 0.01) for each network versus the average of all other network.

In the functional enrichment analysis, to maximize the number of samples, we used data from all six donors (**Figure 5c**) to define the preferential gene sets (**Figure 5d**). While the total number of genes per network increased due to an increase in power, the relative size of each gene set was similar to the gene sets previously defined. The most significantly upregulated genes (FDR < 0.01 for each contrast of network vs. all other networks) were found in the visual network, with 2,243 total preferentially expressed genes. These findings were similar to those in adults which showed the most preferentially expressed genes in the adult limbic and visual networks (Anderson et al., 2018). The default network had the second most preferentially expressed genes, with 1,663 genes. In decreasing order, the number of upregulated genes for the remaining networks were: 409 in the ventral attention network, 287 in the sensorimotor network, and 85 in the high-level vision network (full lists of significantly upregulated genes are found in **Supplementary Data**).

Potential biological and cellular functions and related diseases were identified for each network-specific gene set by comparing the preferentially expressed gene sets to annotated gene sets in all categories through ToppGene’s functional enrichment tool, ToppFun (J. Chen et al., 2009). Full functional enrichment output, including complete lists of genes and all categories, can be found for each network in the **Supplementary Data**. Gene enrichment analyses revealed enrichment for functions and diseases generally related to the nervous system; here we highlight enrichments for the sensorimotor network genes (**Table 1**), but summaries of the functional enrichment analysis for each network are found in **Supplementary Tables 4—7**).

**Table 1.**
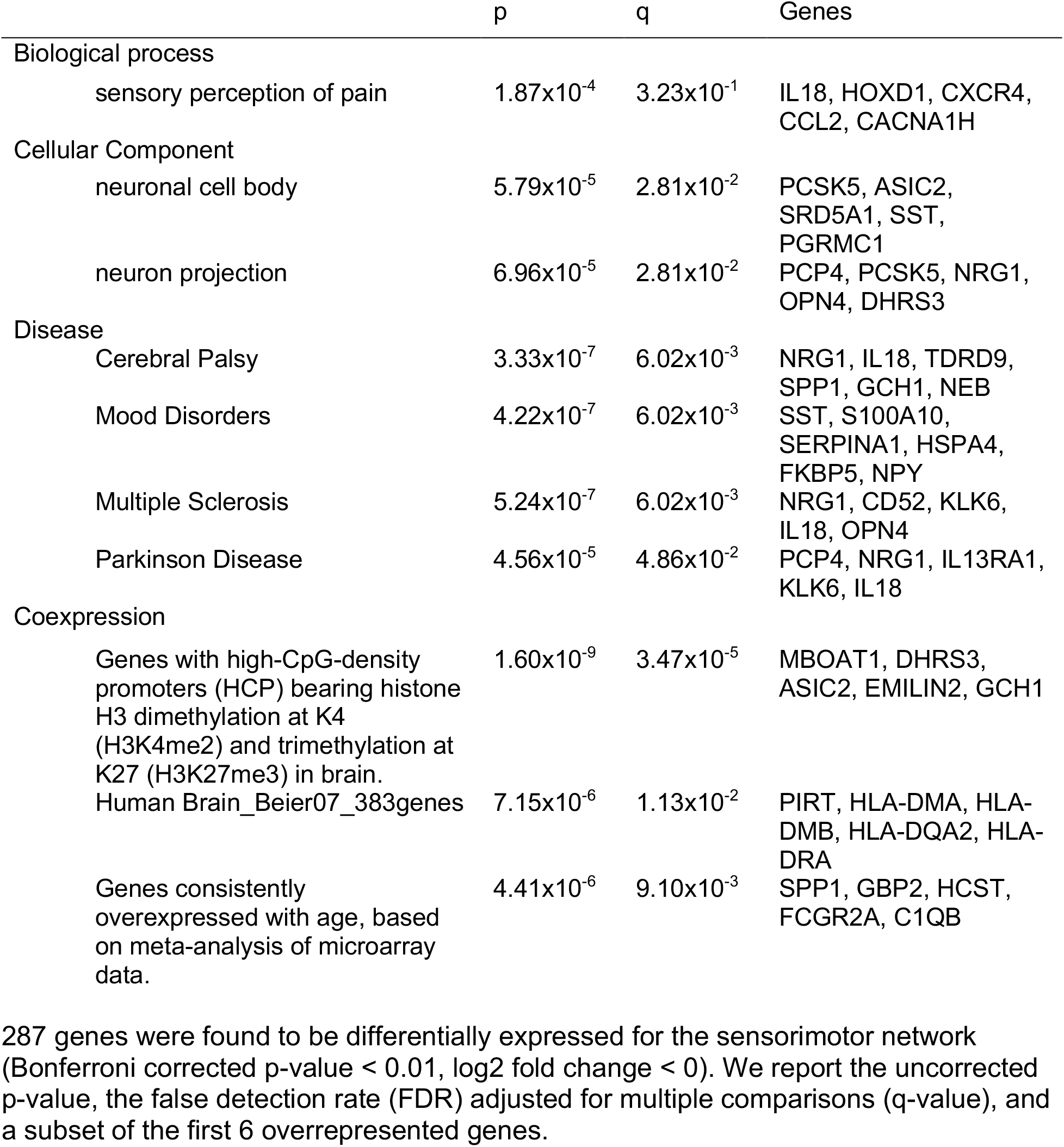
Sensorimotor Enrichment.

|  | p | q | Genes |
| --- | --- | --- | --- |
| Biological process |  |  |  |
| sensory perception of pain | 1.87x10<sup>-4</sup> | 3.23x10<sup>-1</sup> | IL18, HOXD1, CXCR4, CCL2, CACNA1H |
| Cellular Component |  |  |  |
| neuronal cell body | 5.79x10<sup>-5</sup> | 2.81x10<sup>-2</sup> | PCSK5, ASIC2, SRD5A1, SST, PGRMC1 |
| neuron projection | 6.96x10<sup>-5</sup> | 2.81x10<sup>-2</sup> | PCP4, PCSK5, NRG1, OPN4, DHRS3 |
| Disease |  |  |  |
| Cerebral Palsy | 3.33x10<sup>-7</sup> | 6.02x10<sup>-3</sup> | NRG1, IL18, TDRD9, SPP1, GCH1, NEB |
| Mood Disorders | 4.22x10<sup>-7</sup> | 6.02x10<sup>-3</sup> | SST, S100A10, SERPINA1, HSPA4, FKBP5, NPY |
| Multiple Sclerosis | 5.24x10<sup>-7</sup> | 6.02x10<sup>-3</sup> | NRG1, CD52, KLK6, IL18, OPN4 |
| Parkinson Disease | 4.56x10<sup>-5</sup> | 4.86x10<sup>-2</sup> | PCP4, NRG1, IL13RA1, KLK6, IL18 |
| Coexpression |  |  |  |
| Genes with high-CpG-density promoters (HCP) bearing histone H3 dimethylation at K4 (H3K4me2) and trimethylation at K27 (H3K27me3) in brain. | 1.60x10<sup>-9</sup> | 3.47x10<sup>-5</sup> | MBOAT1, DHRS3, ASIC2, EMILIN2, GCH1 |
| Human Brain_Beier07_383genes | 7.15x10<sup>-6</sup> | 1.13x10<sup>-2</sup> | PIRT, HLA-DMA, HLA-DMB, HLA-DQA2, HLA-DRA |
| Genes consistently overexpressed with age, based on meta-analysis of microarray data. | 4.41x10<sup>-6</sup> | 9.10x10<sup>-3</sup> | SPP1, GBP2, HCST, FCGR2A, C1QB |

287 genes were found to be differentially expressed for the sensorimotor network (Bonferroni corrected p-value < 0.01, log2 fold change < 0). We report the uncorrected p-value, the false detection rate (FDR) adjusted for multiple comparisons (q-value), and a subset of the first 6 overrepresented genes.

Genes preferentially expressed in the sensorimotor network were related to particular synaptic and presynaptic neuronal components, but also showed enrichment for certain diseases that affect motor function including cerebral palsy, multiple sclerosis, and Parkinson’s disease (**Table 1**), consistent with previous work supporting the involvement of this network in these disorders (Burton et al., 2009; Rocca et al., 2008; Tessitore et al., 2014; Zhuang et al., 2015).

## Discussion

In adults, brain regions fluctuate with one another at rest, organizing the brain into distinct functional networks that support complex mental functions and vary across individuals. We investigated this spontaneous, intrinsic brain activity in a large cohort of newborns scanned within one week of life and used an entirely data-driven approach to cluster these dense voxelwise connectomes into discrete networks on both group and individual-levels.

We discovered that even in the neonate brain, connectivity profiles of neighboring voxels are distinct enough to be separable into different networks. Much like those observed in adults, neonates had both local and distributed networks that emerged hierarchically, were symmetrical, and, crucially, were replicated across two independent groups. Quantitative comparisons to the adult networks revealed that sensorimotor and visual networks were adult-like, but that frontoparietal cortices were less dissociable into separate networks in neonates, suggesting immature connectivity profiles in these regions. While some previous work is consistent with these findings (Fransson et al., 2007; Gao et al., 2015) but see Doria et al. 2010) our approach provides binary parcellations rather than weighted resting state networks (e.g. through ICA). Critically, because our approach does not rely on assumptions of adult-like network presence in infants, but instead chooses network solutions based on fit and replicability, we found that specifically it is the (frontomedial) control and limbic networks that are immature in neonates. The optimal 5-network solution for neonates did not show these regions as discrete networks. Even when given the opportunity to split into different networks (i.e., at higher clustering solutions), the neonate connectome simply further split up the visual cortices (especially the ventral pathways) rather than splitting these frontomedial cortices into separate networks.

Instead, we discovered the following hierarchy of networks with increasing k-solutions: The first network to dissociate from the rest of the brain was the sensorimotor network which included visual and somatomotor cortex. Next, the visual cortex (both early and late visual cortex) split off from the sensorimotor network, followed by further divisions of visual cortex into a low and high-level visual cortex in the 5-network solution. The high-level visual network clearly displays a split into dorsal and ventral streams, suggesting that perhaps the connectivity of these regions is already set up to perform these high-level functions, which is in line with recent work in infants and neonates (Cabral et al., 2020; Kamps et al., 2020; Li et al., 2020). The visual and high-level visual clusters then further split into anterior/posterior and medial/lateral divisions. High-level visual cortices have gradients or divisions of functional organization based on a combination of retinotopy, animacy, size and other factors in human adults and other animals (Arcaro & Livingstone, 2017; Gomez et al., 2019; Grill-Spector & Weiner, 2014; Hasson et al., 2002; Konkle & Caramazza, 2013; Malach et al., 2002); these gradients and divisions closely mimic the connectivity-based divisions that we observe here in human neonates. The present results suggest that the proto-organization of category-selective visual cortex is not merely retinotopic (not simply clustered within early visual cortex), but rather that it may have important domain-specific connections that are already present at birth.

In contrast to the visual cortices, adult-like control and limbic networks in the frontal cortices were not observed in neonates. The higher clustering solutions that were stable and replicable mainly yielded further subdivision of visual networks. Neonate connectivity did not dissociate between default mode cortex and other frontoparietal and medial frontal cortex.

Because of this, neonates had a single large frontoparietal network (here, called the default mode network due to the high overlap with the Yeo default mode network), which subsumed the area where the control network is in adults. Additionally, the optimal neonate solutions did not have any network which resembled the limbic network in Yeo et al., (2011). The neonatal regions in the proximity of the adult limbic network consisted of multiple, distinct connectivity signatures and were thus part of four different distributed networks (all neonatal networks except for visual). These differences from the adult networks were not attributable to individual variability or within-subject instability. Overall, these results show that domain-specific regions, such as categorical visual areas, have a distinct connectivity signature that separates them from adjacent cortices even at birth, whereas cortices involved in more domain-general processing, such as the cognitive control network, do not have distinct connectivity patterns and may need relevant experience to entrain and preferentially fluctuate with, e.g., other nodes in the control network.

We then identified individual subject solutions, in addition to the group-level network solutions, demonstrating the utility of this approach in exploring individual variability. We found particular networks, such as the ventral attention network, were more variable across individuals despite high within-subject consistency. The sensorimotor and visual areas were the most consistent across individuals, and the ventral attention network had the most between-subject differences, replicating patterns previously found in adults (Kong et al., 2019). This variability is likely not a result of noise, as within-subject network variability was high and consistent across networks. These inter-individual differences may be attributed to differences in developmental trajectory. Previous work exploring group-level networks and cortical growth found that these functional networks and brain regions have different developmental trajectories over the first two years, with sensorimotor and visual areas maturing earlier than attention or control networks (Dehaene-Lambertz & Spelke, 2015; Gao et al., 2015; Gogtay et al., 2004). The degree of individual differences observed here also follows this succession, suggesting individual differences may converge with development and maturation. Alternatively, this individual variability in the ventral attention network may indicate some stable source of individual differences, as the ventral attention network has also been found to exhibit the highest inter-individual variability in adults compared to other networks (Kong et al., 2019). Kong et al. also found that this variability was not due to confounding factors (e.g., network size), but it did relate to individual differences in behavior. Future work can use the connectivity-centroids identified here and apply them to individual subject connectomes (as we do here) to delineate individual solutions for participants and track them across time to better understand the divergence or convergence of individual variability across development and its relationship to individual variation in behavior.

Finally, we showed the dissociability of these networks can be at least partially explained by the underlying genetic expression in these cortices. This result suggests the proto-organization of cortex as outlined here, may be determined in large part by early genetic instructions. The genetic enrichment analyses revealed significant gene sets that are related to nervous system development, and diseases with deficits related to the known function of the network (e.g., preferentially expressed genes in the sensorimotor network were related to cerebral palsy). The differential genetic expression analyses corroborated similar analysis of the adult networks (Anderson et al., 2018). We also observed higher within- than between-network genetic expression and identified similar magnitudes of upregulated genes (Anderson et al., 2018)^32^. These similarities suggest that the neonate networks are also dissociable based on underlying differences in genetic expression.

Future work is necessary to determine the developmental trajectory of these networks (including possible genetic changes and relationship to future behavior) and to address methodological limitations. First, only full-term infants were analyzed here. Identifying networks in preterm neonates, or in longitudinal resting state sessions throughout the first few years of life, could clarify the timeline of the development of these networks and could ascertain if individual differences within networks converge or diverge with development. Second, the networks we have identified are putative functional networks with nomenclature based on topography or overlap with adult networks. Longitudinal studies could identify the relationship between these networks and various mental functions as they manifest, e.g., measured via behavioral measures, such as language and literary skills (Yu et al., 2020), and/or neural measures, such as task-based fMRI (Saygin et al., 2016). Third, the genetic analyses are based on adult tissue samples, and are strictly exploratory. Future analyses can explore the developmental progression of the various preferentially expressed gene sets identified here.

Additionally, as genetic variation affects cortex structure in adults and neonates, further work may reveal if individual variability in functional networks relates to genetic variability. Finally, some methodological differences could affect the comparison with adult networks. The neonate connectome was computed based on voxel to voxel connectivity unlike the Yeo et al. adult connectomes which were computed on a slightly lower resolution supravoxel space. We believe this is actually a strength of the current method, especially because the neonate brain is much smaller than the adult brain. Additionally, while there are differences in the unsupervised clustering algorithms used, the Yeo networks remain relatively stable across parcellation methodologies and datasets (Yeo et al., 2014). Further, our analyses were performed based on currently available, state-of-the-art image processing methods and standard preprocessing techniques. Future improvements of developmental neuroimaging methods will likely improve the results and inferences drawn from the current study.

We provide a way for researchers to apply our clustering solutions to their individual subject connectivity data such that they can obtain subject-specific networks for their data (i.e. by identifying which cluster best matches each voxel’s connectivity pattern; available here: https://github.com/SayginLab/neonate_molloy). These subject-specific networks address a growing need for more personalized models in predicting future outcomes, as discussed in Y. Chen et al., (2021). We also provide the atlas for each k clustering solution so that the clusters can simply be overlaid on subject or group data for e.g. seeds and targets for connectivity analyses or as regions of interest for fMRI analyses. Overall, we identified underlying neonatal networks from resting state data that can be identified on an individual level and are explained by genetic data. These findings indicate some proto-organization at birth, while also highlighting the necessity of experience for full adult-like organization.

## Methods

### Neonate Dataset

The neonate data are a part of the Developing Human Connectome project (Hughes et al., 2017), a large-scale study of the structural and functional organization of the developing brain. The resting state data comprises of a single 15 min scan, and all of the included infants were scanned during natural sleep. The Developing Human Connectome project was approved by the UK Health Research Authority (Research Ethics Committee reference number: 14/LO/1169). All infants were recruited and scanned at the Evelina Newborn Imaging Centre, St Thomas’ Hospital, London, UK and written parental consent was obtained for each individual in the release. The second release consists of 505 subjects. Only full-term neonates (born at greater than or equal to 37 gestational weeks) with at least one session of functional data were included. Individual sessions were included if there was no sedation during a session, successful group registration (missing less than 10 voxels in registration to template), and had radiological scores corresponding to “normal” findings, or findings with unlikely clinical/ analysis effects. The resulting data consists of 267 full term neonates (121 females, 146 males). The average age at scan was 1.3 chronological weeks (mean gestational age at scan = 41.10 weeks, see full distributions in **Supplementary Figure 1**).

### Acquisition Details

Neonates were scanned using a 3T Philips Achieva (modified R3.2.2 software) with a dedicated neonatal imaging system containing a 32 channel phased head coil (Hughes et al., 2017). High-resolution anatomical images (T2-weighted and inversion recovery T1-weighted multi-slice fast spin-echo images) were acquired with in-plane resolution 0.8x0.8mm^2^ and 1.6mm slices overlapped by 0.8mm (T2w: TE/TR = 156/12000 ms; T1w: TE/TR/TI = 8.7/4795/1740 ms). Resting-state fMRI data using multiband (MB) 9 × accelerated echo-planar imaging for neonates during natural sleep (sedated infants excluded) were collected (TE/TR = 38/392 ms, voxel size = 2.15 × 2.15 × 2.15 mm^3^) for a total of approximately 15 min (2300 volumes). Single-band reference scans with bandwidth matched readout, and additional spin-echo acquisitions with both AP/PA fold-over encoding directions were also acquired.

### Preprocessing

The functional data were processed following the dHCP preprocessing pipeline (Fitzgibbon et al., 2020; Makropoulos et al., 2018) and were registered to the 40-week template (Schuh et al., 2018). Additional details summarizing the minimally preprocessing steps are found in the **Supplementary Materials** and examples of subjects not successfully registered to the group template are shown in **Supplementary Figure 2**. Only the cortical gray matter, deep subcortical gray matter, and hippocampus/amygdala voxels were included for connectivity analysis and these were defined according to tissue segmentation masks from the dHCP supplied group atlas (Schuh et al., 2018).

### Defining the Connectome

The 267 neonates were then split into a training set of 134 individuals used for tuning the clustering algorithm and a test set of 133 individuals to assess generalizability and replicability (see **Supplementary Table 3** for a summary of sex and age in the training and test set). The connectome was defined as the Pearson correlation between each voxel’s normalized time series at rest. For the group-average connectomes, individual connectomes were transformed using a Fisher’s r to z transform, averaged across the group (either test or train), then transformed back using the inverse Fishers z to r transform. We calculated the connectome for both the whole brain (i.e., cortex and subcortex, resulting in a 23,841 by 23,841 voxel matrix) as well as just the cortex (20,130 by 20,130 voxel matrix) for the training and test sets. We report results from the cortical solutions in the main text and whole-brain solutions in **Supplementary Figure 5**.

### Defining Networks

We used k-means clustering to define networks on the connectome matrices. K-means clustering is an exploratory, unsupervised learning approach that reveals patterns in the underlying data based on the similarities of features in the observations, which in our case is the similarity of the functional connectivity vectors for each voxel. Similarity was quantified using correlation distance between connectivity vectors; correlation distance captures similarities in connectivity profiles, as opposed to other methods which may capture similarities in the magnitude of connectivity (driven by differences in signal-to-noise ratio, see **Supplementary Figure 4**). Further, this metric is equivalent to the distance metric used by Yeo et al. Five replicates were specified to ameliorate the effect of initial values. The resulting networks were color-coded for illustrations based roughly on the coloring scheme of the 7-network solution in Yeo et al..

Because the k-means clustering algorithm is dependent on our choice of the underlying number of clusters, or k, we calculated solutions along a sequence of k clusters ranging from k= 2 to 25. These solutions were independently calculated for training group average connectome and the testing group average connectome and evaluated based on fit and stability. The results for all cortical test and train solutions across this entire sequence are shown in **Supplementary Figures 6—10**.

An optimal clustering solution should fit the training data well, but also should generalize to the testing data. To fit the data well, a solution should maximize the similarity between voxels in the same cluster and minimize similarity between voxels in different clusters (i.e., maximize separation). We quantified this ‘fit’ using a metric based off of the silhouette coefficient and functional homogeneity (Shi et al., 2018). A silhouette coefficient is calculated for every cluster, *k*, in a given solution. The functional homogeneity silhouette coefficient is defined as

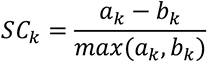

where *a_k_* is the functional homogeneity within cluster *k* and *b_k_* is the functional homogeneity between voxels in cluster *k* and other clusters. The within cluster functional homogeneity, *a_k_* is defined as

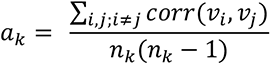

where *n_k_* denotes the number of voxels in cluster k, *corr*(*v_i_*,*v_j_*) is the correlation between voxel *i* and voxel *j* (i.e., functional homogeneity of the average training data), and *i* and *j* range from 1 to *n_k_*. The between cluster functional homogeneity, *b_k_*, is defined as

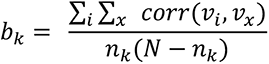

where *n_k_*again denotes the number of voxels in cluster k, *corr*(*v_i_*,*v_x_*) is the correlation between voxel *i* in cluster k and voxel *x* outside of cluster k (i.e., functional homogeneity of the average training data), *i* ranges from 1 to *n_k_* (including all voxels within cluster *k*), and *x* ranges from 1 to *N* − *n_k_*, where N is the total number of voxels (in other words, including all voxels not in cluster *k*). An ideal solution would maximize *a_k_* and minimize *b_k_*, so an ideal solution would have a larger *SC_k_*. The possible range of *SC_k_* is -1 to 1, where 1 indicates the best possible solution, and -1 indicates the worst possible solution.

Additionally, a solution should generalize to other data well (here, the testing data), to avoid overfitting. A generalizable solution would have a high degree of overlap between the training solution and the testing solution. We use two metrics to generalize overlap: the dice coefficient (Dice, 1945) and the Adjusted Rand Index (ARI; Vinh et al., 2009)).

The dice coefficient has been widely used in studies comparing brain parcellations in adults, and we use the same measures used in Arslan et al.,(2018). Dice ranges from 0 to 1, where 1 indicates perfect overlap and is defined as

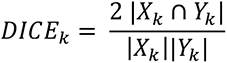

Where |*X_k_* ∩ *Y_k_*| is the number of voxels in both parcel X and parcel Y, |X_k_| is the total voxels in X_k_, and |Y_k_| is the total voxels in Y. Dice is computed for pairs of parcels in two different solutions (here, test and train) with the largest overlap (without replacement), so there is a dice value for each cluster in a given solution. To choose our optimal solution, we use the mean dice.

The ARI is another measure of overlap, but unlike dice, it does not require the pairing of parcels. We use ARI as defined in (Arslan et al., 2018) as well. The ARI for two solutions X and Y is

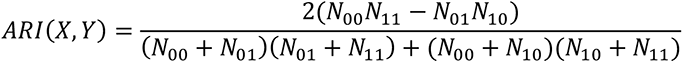

where *N*_11_ indicates the number of pairs of voxels that are assigned to the same cluster in both solution X and solution Y, *N*_00_ indicates the number of pairs of voxels that are assigned to different clusters in both solution X and solution Y, *N*_01_ indicates the number of pairs of voxels that are assigned to different clusters in solution X but the same cluster in solution Y, and *N*_10_ indicates the number of pairs of voxels that are assigned to the same cluster in solution X, but different clusters in solution Y.

To define symmetry, we quantify the proportion of voxels with the same network assignment on both hemispheres. In the neonate group template, each voxel in the right hemisphere is projected to the left hemisphere and matched to the nearest left hemispheric voxel. If the left and right voxels have the same network assignment, it is considered symmetric.

The overall symmetry measure for a given solution is the proportion of matches for all right hemispheric voxels. Symmetry broken up by network is reported in **Supplementary Figure 13**. **Individual Solutions**

We used the group centroids that were calculated from the training set as our prior, and then ran a k-nearest neighbor classifier to assign voxels to the most similar connectivity profile. We did this for every individual in the test set, so we had a total of 133 individual solutions.

Between-subject overlap was quantified per voxel by calculating the proportion of individuals with the same network assignment for that voxel (using the independent training set as ground truth for network assignment). Average overlap (using the dice coefficient), and the standard error of the mean across all voxels within each network was also calculated. To calculate within-subject stability of each network, we split the 15-minute resting state scan into 5-minute segments, then concatenated those sections into three 10-minute blocks, calculated the connectome, and calculated the individual solutions. Then, we quantified the within-subject overlap by finding the dice coefficient of overlap across those three solutions. The within- and between-subject overlap for each network was compared using (paired, non-parametric, two-sided) Wilcoxon signed-rank tests. Two additional tests compared the ventral attention network vs. the average overlap of all other networks for within and between subject overlaps.

### Comparison to Adult Networks

The comparison of infant and adult networks was done in (volumetric) FreeSurfer CVS average-35 MNI152 space. The neonate group solutions were linearly transformed (with nearest neighbor interpolation) from the infant template Schuh et al. space to the adult MNI space (**Supplementary Figure 3**). To account for differences in coverage and fill in gaps from the registrations, the infant solutions were dilated 4 times with a 6 mm kernel, then non-gray matter cortical voxels were removed (defined by tissue masks). The adult 7-network Yeo (liberal mask) solution in MNI space was downloaded from https://surfer.nmr.mgh.harvard.edu/fswiki/CorticalParcellation_Yeo2011. Note that the Yeo et al. solution was computed on the surface, but because of the lack of robust surface methods for neonates, we complete all analyses in the volume space. We calculated the overlap between adult and neonate (the testing solutions for k = 5 and k = 7) solutions using the dice coefficient as outlined above. Only voxels present in both the MNI images of the Yeo and neonatal solutions were compared.

### Genetic Expression by Network

Normalized microarray data were obtained from the Allen Human Brain Atlas (AHBA; Hawrylycz et al., 2012). These data include samples of 20,787 genes from 6 donors. Four donors contained only left hemispheric samples, while the other 2 contained bihemispheric samples. To explore within/between network differences in genetic expression, the data from the four left hemispheric donors defined the network-specific gene sets, and the data from the bihemispheric donors were used to compare the mean expression of those gene sets within the relevant network and between all other networks (following similar methods as outlined in Anderson et al., 2018 for adult networks). We included data from all 6 donors to determine each network’s preferential genes and enrichment. The microarray data were further preprocessed following steps from (Gomez et al., 2019). First, we excluded probes without a gene symbol or Entrez ID. Second, we averaged the expression profiles of all probes targeting the same genes (within an individual). Third, we standardized within each donor.

Before matching the AHBA samples to the neonatal networks, we registered both the AHBA samples and the neonatal networks to FreeSurfer CVS average-35 MNI152 space (details in **Supplementary Materials**). Additionally, we split the large neonatal networks into smaller subsections. First, we separated discontinuous components using nilearn (Pedregosa et al., 2011), separating by hemisphere and excluding regions less than 100 mm^3^. Second, very large subregions (more than 1,000 voxels) were further divided using a watershed algorithm (Meyer, 1994). Finally, the watershed-defined regions were slightly dilated within the boundaries of the original networks to define the final subregions to which the AHBA samples were matched. Additional matching details including number of subregions matched per individual and network are included in the **Supplementary Table 3.** Expression values used in the genetic analyses averaged individual samples within the same subregion.

Preferentially expressed genes for each network were compared to the expression of that gene in the average of all other networks using a linear model in the limma package in R (Ritchie et al., 2015). We accounted for subject-to-subject (across donor) differences using the donor as a blocking variable and the residual donor effects were controlled for using limma’s duplicateCorrelation tool. Genes were identified as significantly upregulated only if they passed Bonferroni correction at q ≤ 0.01 and log2 fold-change (logFC) > 0. Using the samples from the four left-hemispheric donors as the test set, we identified the preferential genes for the five networks. We found a total of 2,937 unique preferential genes for the five networks. The gene set for the visual network was the largest (1,638 genes), but preferentially expressed genes were identified for all networks (1,000 default genes, 103 ventral attention genes, 179 sensorimotor genes, and 12 high-level vision genes). The mean expression for each network’s set of genes was calculated for each sample in the test set (remaining two donors), resulting in both within-network and between-network mean expression observations for each sample. We used an ANOVA model (fixed effects for network expression, random effect of donor) to identify if the within-network expression was higher than the between- network expression. Because these two left out donors contain samples from both hemispheres, the total number of matched observations (465) was comparable to the total number of observations used to define the gene sets (456). Post-hoc tests (one-way ANOVA, random effect of donor) revealed the degree of within- versus between-network differences. We report p-values, F-statistics, effect sizes as indices of partial variance explained (η_p_^2^), and confidence intervals of the estimated marginal mean difference of between expression – within expression, using the Satterthwaite method.

For reporting the preferentially expressed genes and the enrichment analyses, we used data from all six donors, performing an identical limma model analysis as above. Preferentially expressed genes were again defined as those genes with positive logFC and that passed Bonferroni correction at p < 0.01 for each contrast/network. The gene enrichment analyses were performed with ToppGene Suite (J. Chen et al., 2009). The enrichment gene sets were defined using the Entrez IDs of the genes identified by each network’s preferentially expressed genes. All features were included, with a FDR corrected p-value cutoff of 0.05.

## Acknowledgements

Data were provided by the developing Human Connectome Project, KCL-Imperial-Oxford Consortium funded by the European Research Council under the European Union Seventh Framework Programme (FP/2007-2013) / ERC Grant Agreement no. [319456]. We are grateful to the families who generously supported this work. Analyses were completed using the Ohio Supercomputer (https://www.osc.edu). This research was partly funded by the Alfred P. Sloan Foundation (to Z.M.S), Ohio State’s Chronic Brain Injury Program (to Z.M.S), and Ohio State University’s College of Arts and Sciences (to Z.M.S. and M.F.M.).

## Data availability

The neuroimaging data used in this study are available as part of the publicly available developing Human Connectome Project (dHCP; https://www.developingconnectome.org). The genetic data are from the publicly available Allen Human Brain Atlas (AHBA; http://human.brain-map.org/static/download). Resulting neonate parcellations and full genetic expression results can be found at https://github.com/SayginLab/neonate_molloy.

## Code Availability

Analyses were completed using MATLAB R2020a (The MathWorks Inc., Natick, USA) and R version 4.0.3 (2020-10-10). MATLAB code to calculate individual parcellations will be publicly available at https://github.com/SayginLab/neonate_molloy. All code is available from the authors upon request.

