## Supplementary Materials for "Individual Variability in the Innate Functional Organization of the Human Brain" (from another version of this preprint: version 1, 2021-03-24)

**Supplementary Information for *Innate Organization of the Human Brain***

### Non-imaging Details from dHCP

The second release of the dHCP included a total of 506 subjects, but 42 of those subjects had no functional scans. 416 of the remaining subjects had one functional scan and 48 completed two functional scans. A total of 267 neonates (146 males, 121 females) met the inclusion criteria: 1) at least one session functional data, 2) only sessions with radiological scores corresponding to "normal" findings or findings with unlikely clinical/ analysis effect), 3) no sedation during a session, 4) missing less than 10 voxels in registration to template (see below), and 5) Born at ≥ 37 gestational weeks. The mean gestational age at birth was 39.97 weeks, the mean gestational age at scan was 41.10 weeks, and the mean chronological age at scan was 1.31 weeks (distributions of these variables in **Supplementary Figure 1**). The neonates were then randomly split into two groups, the training group with N = 144 individuals and the test group with N = 143 individuals (summary of sex and age in **Supplementary Table 1**).

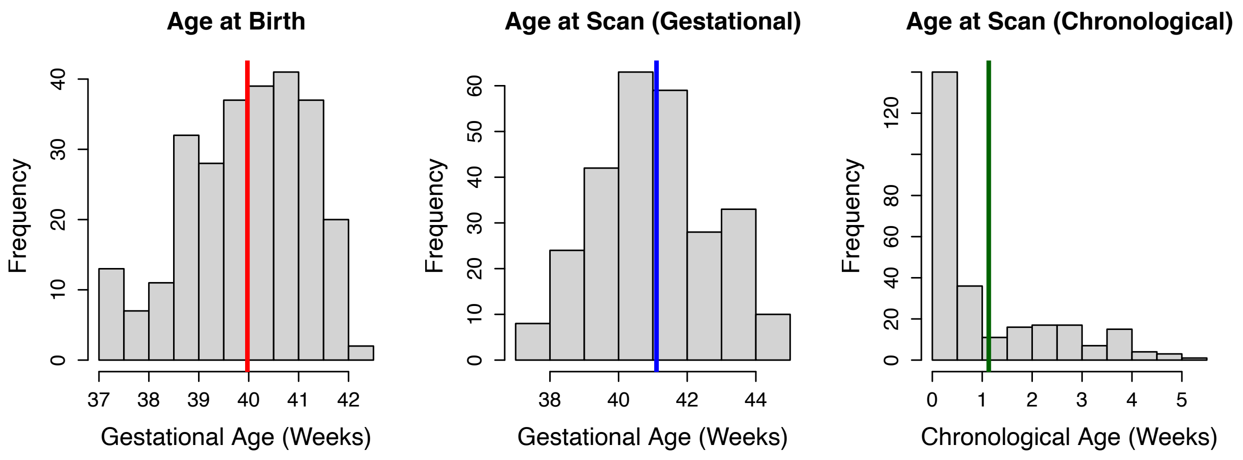


**Supplementary Figure 1**. Distributions of the ages at birth and scan for the 267 included subjects. Vertical lines denote the mean.

**Supplementary Table 1**
***Training and Test Set Non-Imaging Summary***

| | Train (N = 144) | Test (N = 143) |
|---|---|---|
| Sex (Count) | | |
| Female | 67 | 54 |
| Male | 67 | 79 |
| Gestational age at birth (in weeks) | | |
| Mean | 39.82 | 40.13 |
| Standard deviation | 1.24 | 1.15 |
| Gestational age at scan (in weeks) | | |
| Mean | 40.86 | 41.23 |
| Standard deviation | 1.70 | 1.73 |
| Chronological age at scan (in weeks) | | |
| Mean | 1.15 | 1.11 |
| Standard deviation | 1.30 | 1.24 |

**Functional Data Registration**

We use the minimally preprocessed functional data outlined in Fitzgibbon et al. (2020). To summarize these steps in more detail, first, the fieldmap was preprocessed by aligning the susceptibility distortion field with the functional data. Then, susceptibility and motion correction consisted of slice to volume correction, susceptibility distortion correction, and regressing out the motion estimates. Finally, the functional data was denoised by regressing out other nuisance estimates. In addition to the steps outlined by Fitzgibbon et al., we registered the functional images to a standard template space. The images were aligned with the individual native T2 image and the volumetric by nonlinearly mapping the 4D functional image to the 2x2x2 mm 40-week atlas (Schuh et al., 2018) in FSL using the warping information included in the second release. The results of applying the warp to the preprocessed functional data is shown for a single subject below. Note the lack of coverage in posterior cerebellum of this subject was also present in the unprocessed data. This lack of coverage was also used in our decision to mask out the cerebellum for the functional data used to calculate the connectome. We then masked only the areas of interest in our parcellation using the template tissue segmentations from the 40-week template from Schuh et al. (2018). The cortical gray matter, deep gray matter, and hippocampus and amygdala are all included in our parcellation. This then excludes the cerebral spinal fluid, white matter, background (non-brain), ventricles, cerebellum, and brainstem.

The template mask including the areas of interest consisted of 23,841 total voxels. However, post-registration, we found that 52 out of 386 subjects (with all the inclusion criteria except for the missing voxel criteria outlined above) were missing at least 1 voxel. The maximum number of missing voxels was from subject CC00109XX10, with session ses-36901, which is missing 189 voxels from the template. The missing voxels tended to be on the dorsal most parts of the brain, see examples of two excluded individuals (more than 10 missing voxels) in **Supplementary Figure 2**. The first example demonstrates the most extreme case, while the second is a more representative excluded subject.

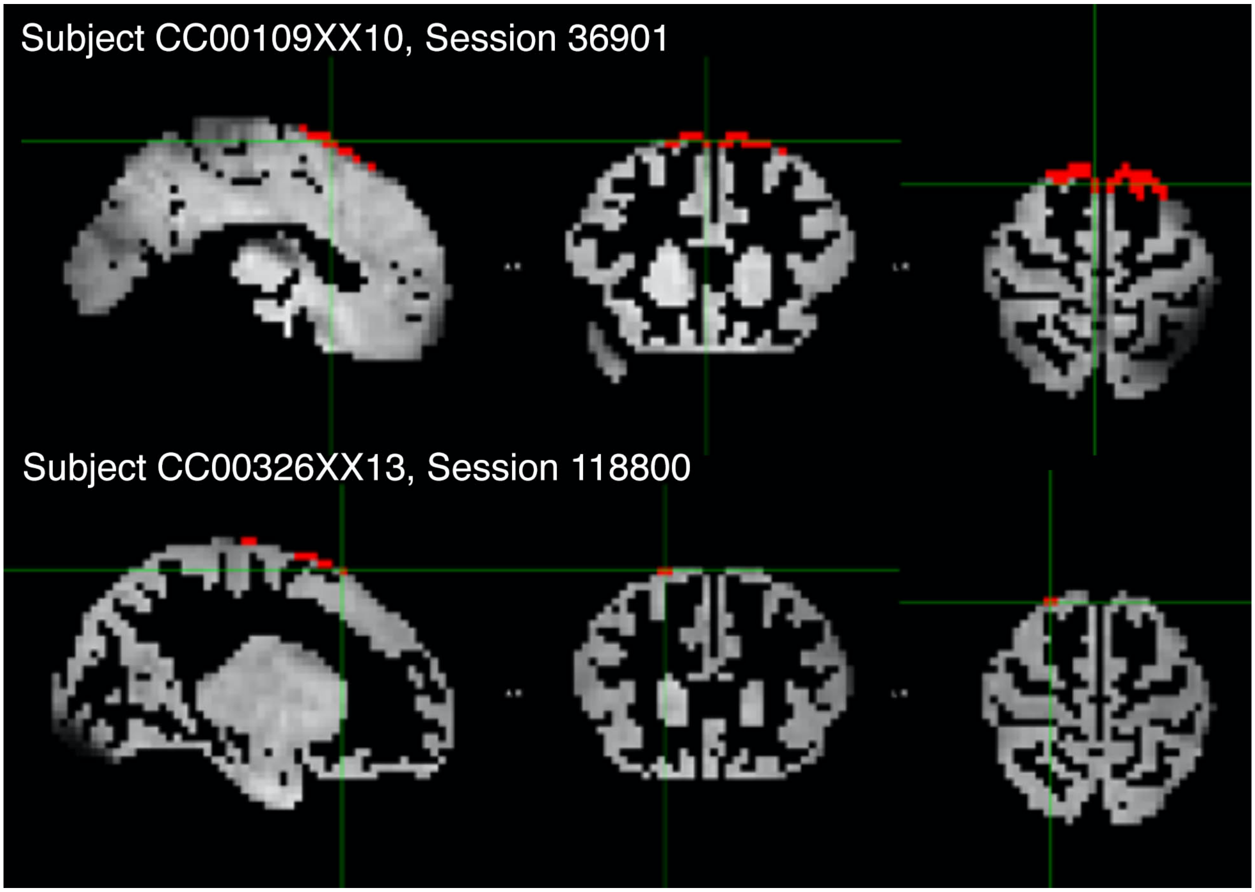

**Supplementary Figure 2**. The mean warped and masked image for the specified individual and session is in grayscale and the template mask of gray matter is red. Red areas indicate that those voxels for that individual and session were not successfully registered to the group template.

For many analyses, including the comparison to adult networks and the genetic analyses, and in order to project to the surface, the neonate results (networks) were linearly registered (using FSL's FLIRT) to the FreeSurfer CVS average-35 MNI152 space (i.e. space that the adult networks were already in). Note that we also attempted to use the infantfs (FreeSurfer) pipeline (Zöllei et al., 2020) but ultimately decided on FLIRT because of better registration. **Supplementary Figure 3** demonstrates that the transformation from infant to MNI space was successful.

### Alternative Distance Metrics

We explored three different metrics: correlation, Euclidean, and cosine. We report only the correlation solutions in the main text for three reasons. First, it is the analogous distance measure used in Yeo et al., (2011) which allows for a more direct comparison to the adult results. Second, the resulting parcellations from the correlation and cosine measures are highly similar. Third, the correlation distance measure captures differences in connectivity profiles, as opposed the Euclidean distance measure which captures signal strength (**Supplementary Figure 4**).

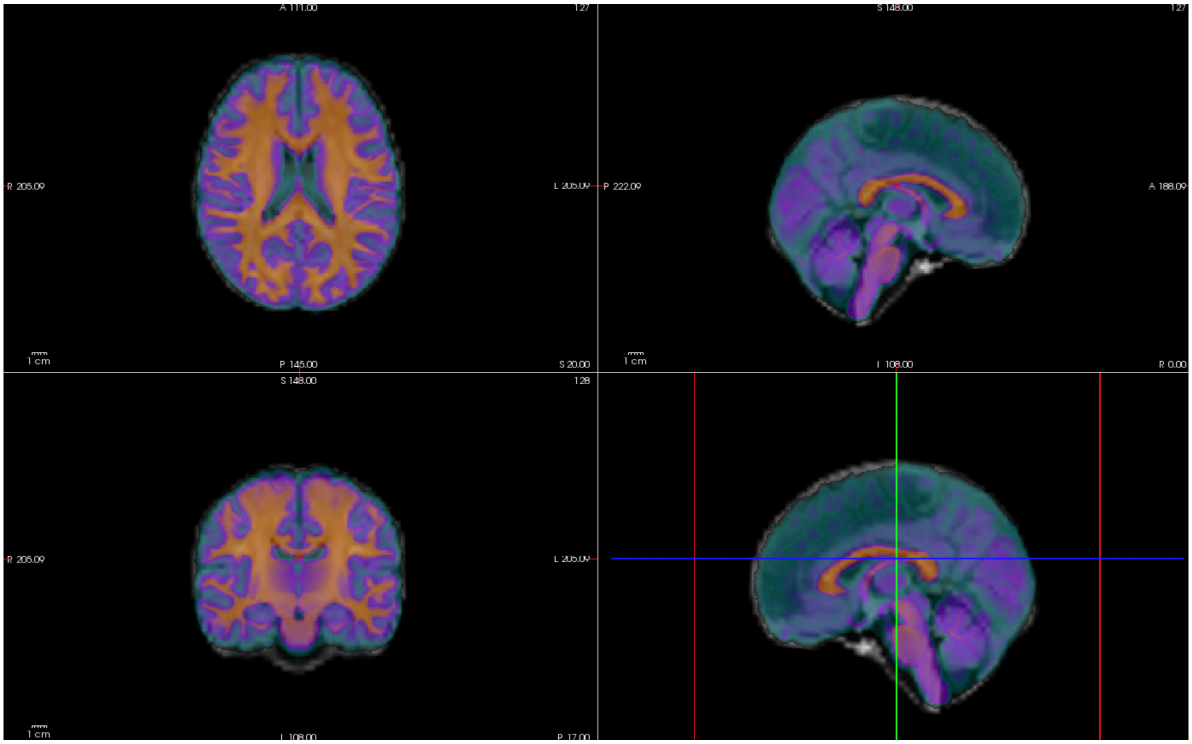

**Supplementary Figure 3**. The grayscale background image is the FreeSurfer CVS average-35 MNI152 anatomical image. The semi-transparent colorful image in the Schuh et al., (2018) 40-week template image after the linear registration to the adult MNI space.

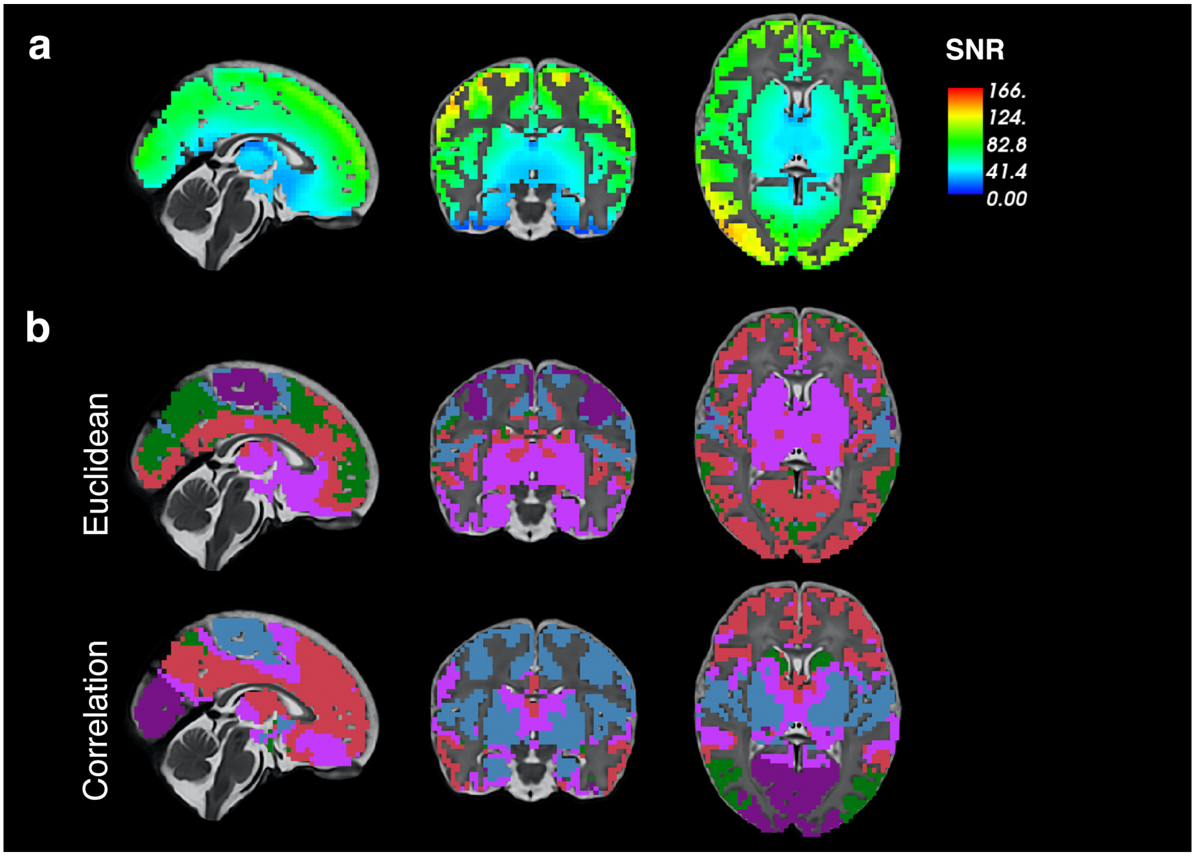


**Supplementary Figure 4**. **a**. Signal to noise (SNR) ratio for each voxel, averaged across subjects in the training set. For each voxel in each subject, the SNR is the mean signal of the time series divided by the standard deviation of that time series. Warmer colors indicate a higher SNR whereas cooler colors indicate a lower SNR. **b**. The k = 5 network solutions of the same underlying training data when specifying the squared Euclidean distance (top row) versus the correlation distance (bottom row). Note that the network borders in the Euclidean solution resemble differences in SNR.

**Whole-brain Optimal Solution**

In additional to the cortical-only solutions presented in the main text, we also found optimal solutions including subcortical voxels. After similarly calculating the fit to data (functional homogeneity silhouette index) and generalizability (test/train overlap), we determined that the k = 5 solution is optimal for the whole-brain solution as well (**Supplementary Figure 5a**). Higher k solutions produced networks that were not reproducible in the testing dataset (contained at least one dice coefficient of 0). **Supplementary Figure 5b** shows the whole-brain 5 network solutions plotted in the volume space for the training and test set. Because the networks closely resembled those in the cortex-only solutions, the naming convention and colors are the same.

**All Cortex-Only Solutions**

The cortex-only solutions for every k calculated from 2 to 25 are shown in **Supplementary Figures 6—10**. All solutions are calculated in the volume space and projected to the surface. The color scheme for each test solution is determined by first finding the overlapping networks (pairwise, from dice coefficients) in the previous solution (e.g., for k = 17, first matching to k = 16) for consistency across solutions. The training solution color scheme is determined by pairwise matches to the test solution for the same k.

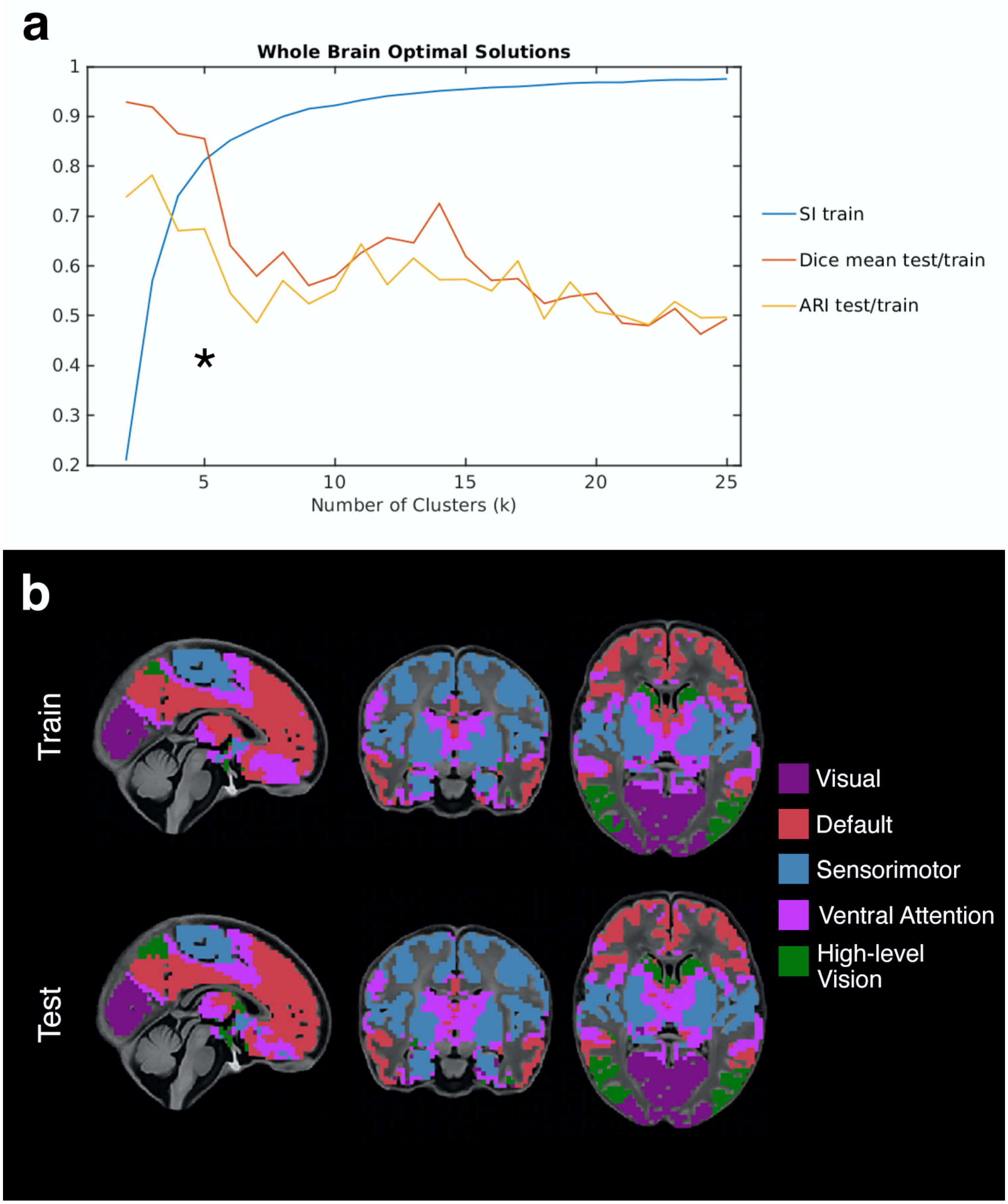


**Supplementary Figure 5. a**. Different quantitative evaluations (y-axis) of the whole-brain clustering solutions for each of the k-clusters (x-axis). The blue line shows the Silhouette Index (SI) for clustering in the training set (essentially the fit of the clustering solutions to the underlying connectivity data). The red and yellow lines show the overlap of the training and test set solutions quantified by the Dice coefficient (red line) and Adjusted Rand Index (ARI; yellow line). **b**. Optimal k = 5 solution for the training and test sets.

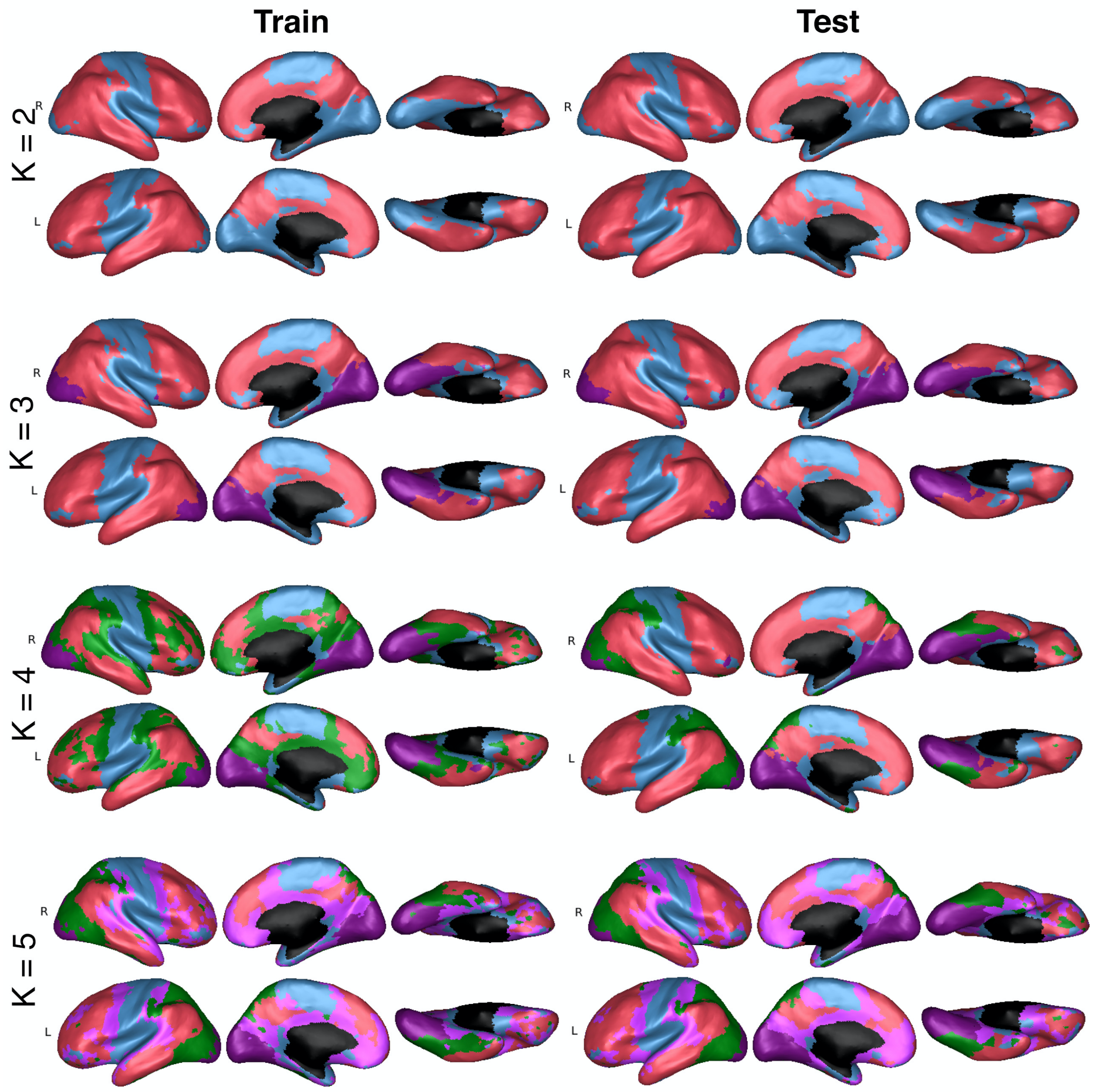


**Supplementary Figure 6**. Training set solutions (left) and test set solutions (right) for k = 2—5 (rows).

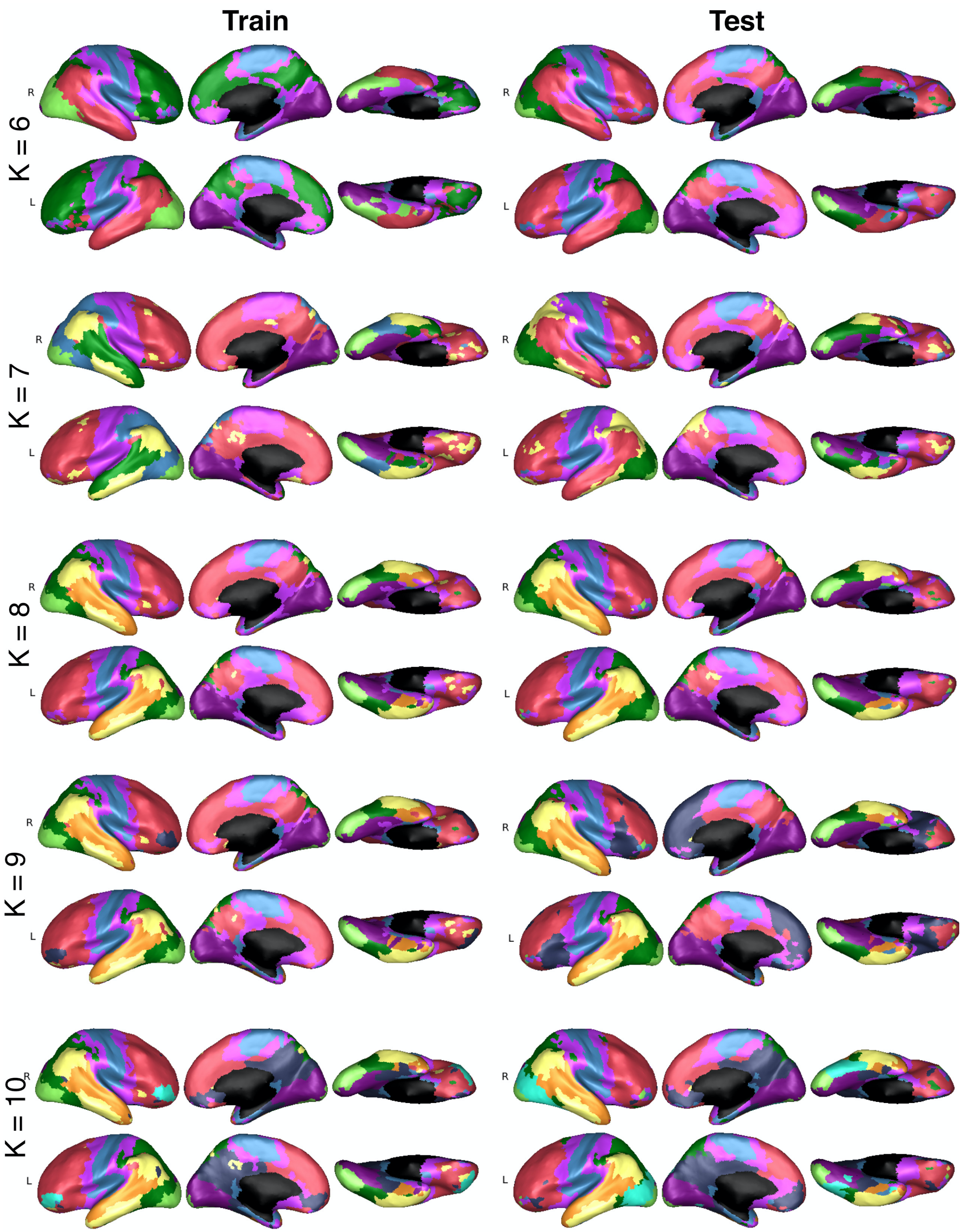


**Supplementary Figure 7.** Training set solutions (left) and test set solutions (right) for k = 6—10 (rows).

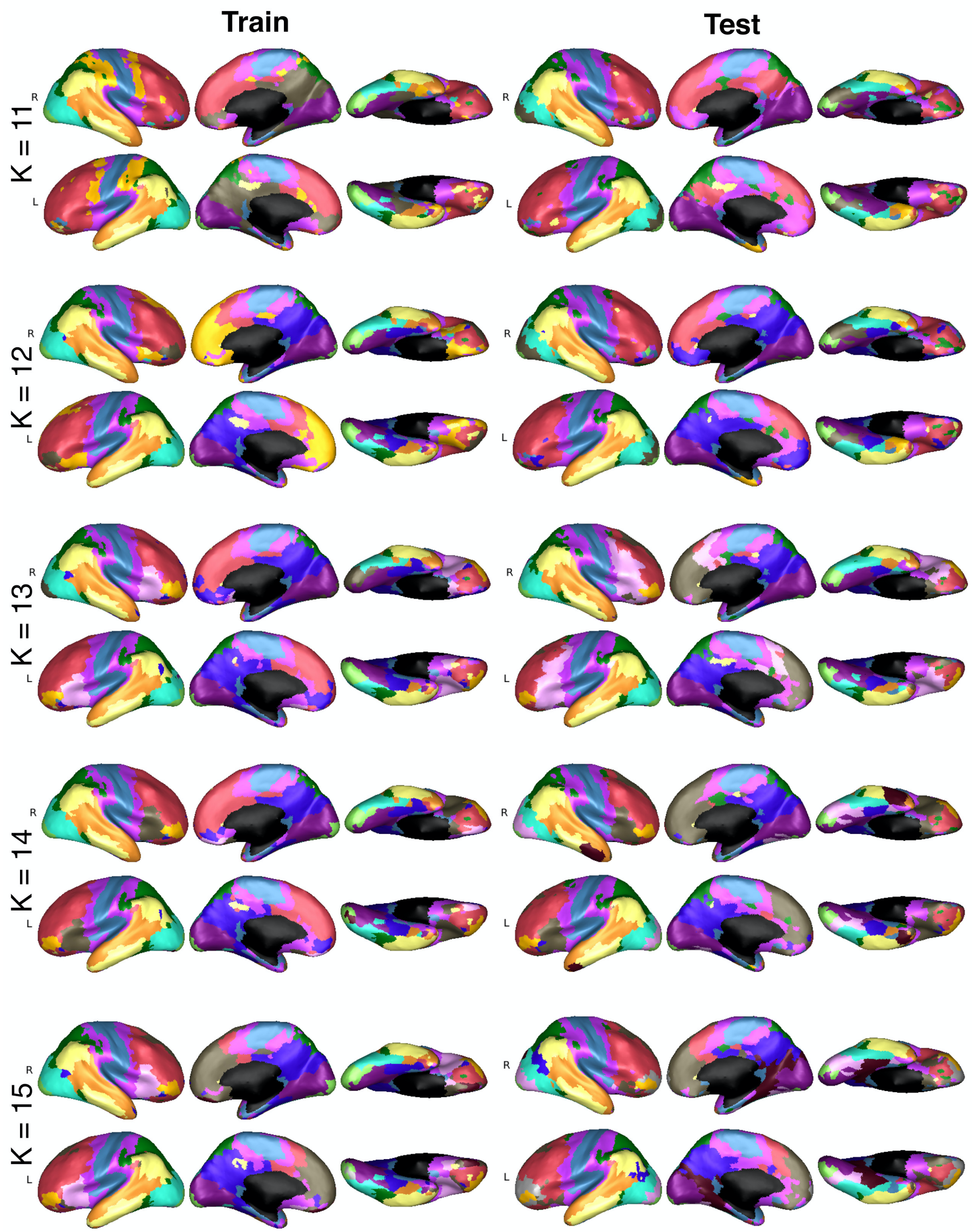


**Supplementary Figure 8.** Training set solutions (left) and test set solutions (right) for k = 11—15 (rows).

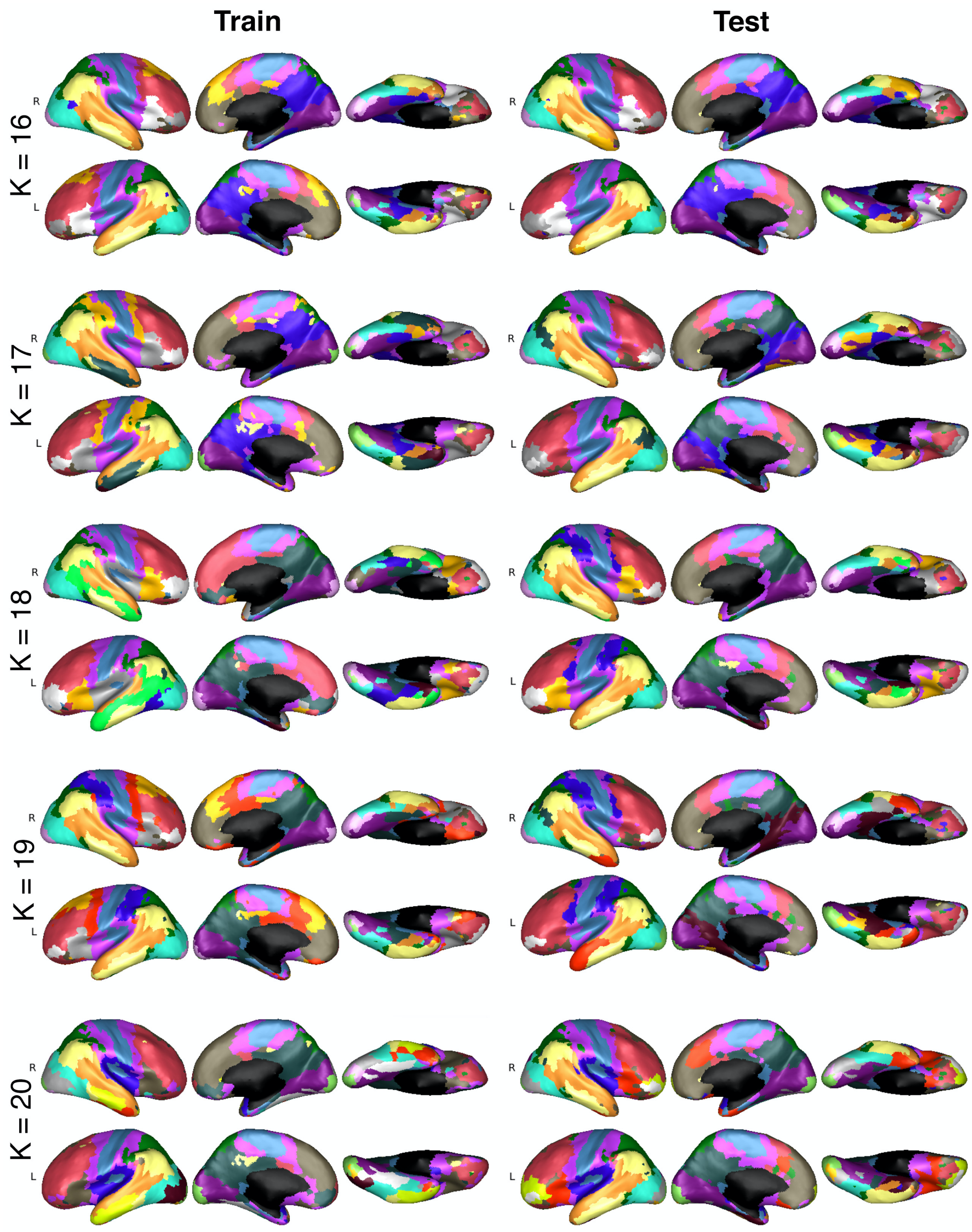


**Supplementary Figure 9.** Training set solutions (left) and test set solutions (right) for k = 16—20 (rows).

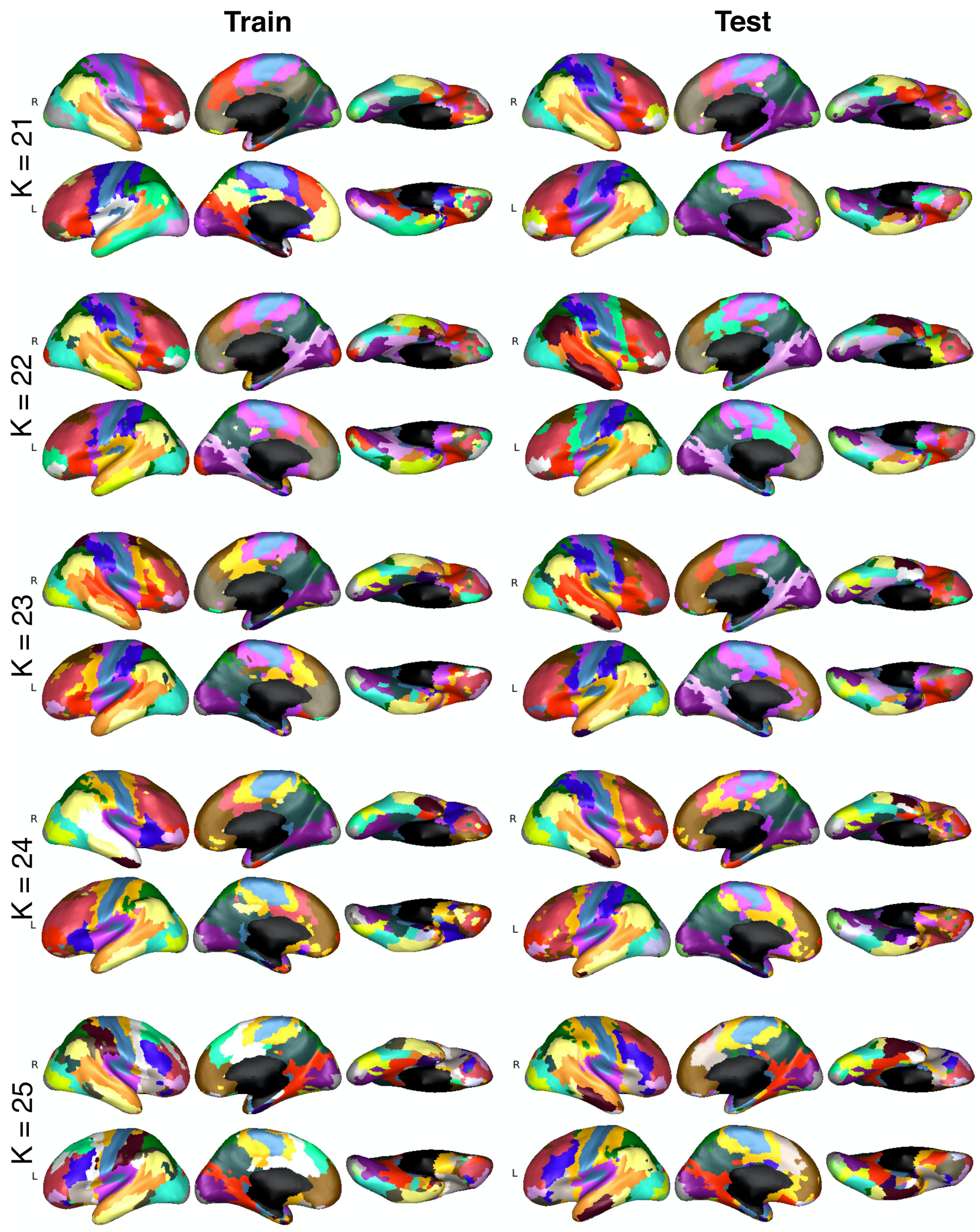


**Supplementary Figure 10.** Training set solutions (left) and test set solutions (right) for k = 21—25 (rows).

### Hierarchical Emergence of Networks

In the independently calculated solutions for different prespecified total networks (k), we observed that the boundaries between networks remained overall stable with increasing networks. Higher resolution networks were then subdivided within one network in the lower resolution solution. This hierarchical emergence of networks was observed in both the whole-brain solutions (**Supplementary Figure 11**) and in the cortical-only solutions (**Supplementary Figure 12, Supplementary Video 1**). Note that each k solution is generated independently from other k solutions, but there is still a clear hierarchy of networks, with initial dissociation of the sensorimotor network from the rest of cortex (blue vs. red), followed by emergence of visual network (purple), then splitting the visual network into low and high-level visual networks (green), then separation of default and ventral attention networks (red and purple).

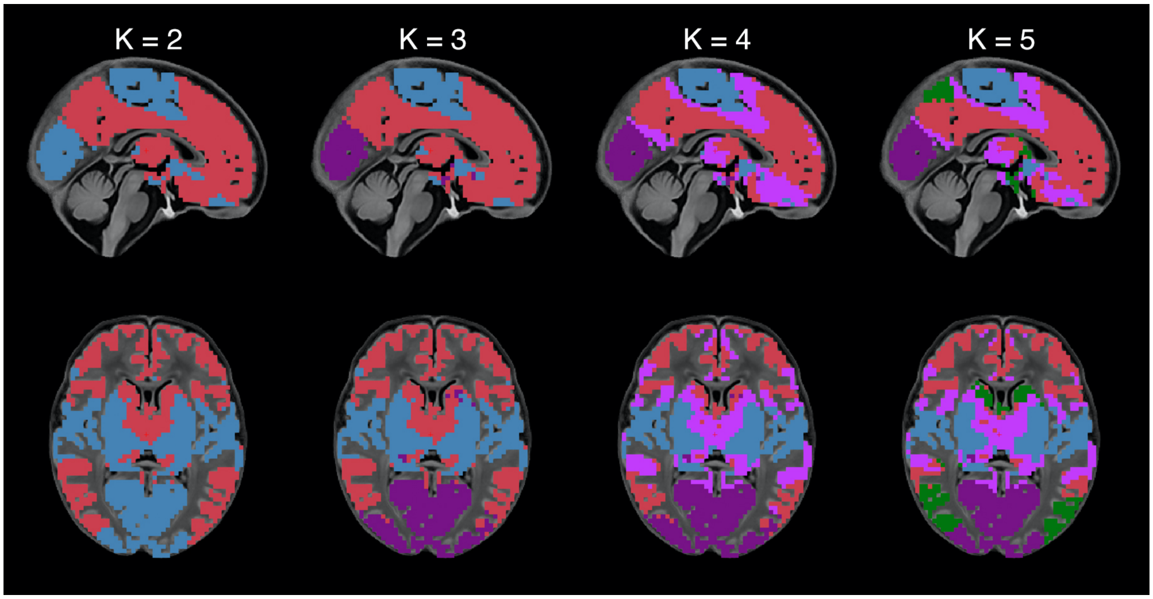


**Supplementary Figure 11.** Each column shows the whole-brain clustering solutions (on the test set of subjects) for an increasing number of clusters (k). Voxels are assigned a k-value based only on the similarity of their connectivity profile to all other voxels

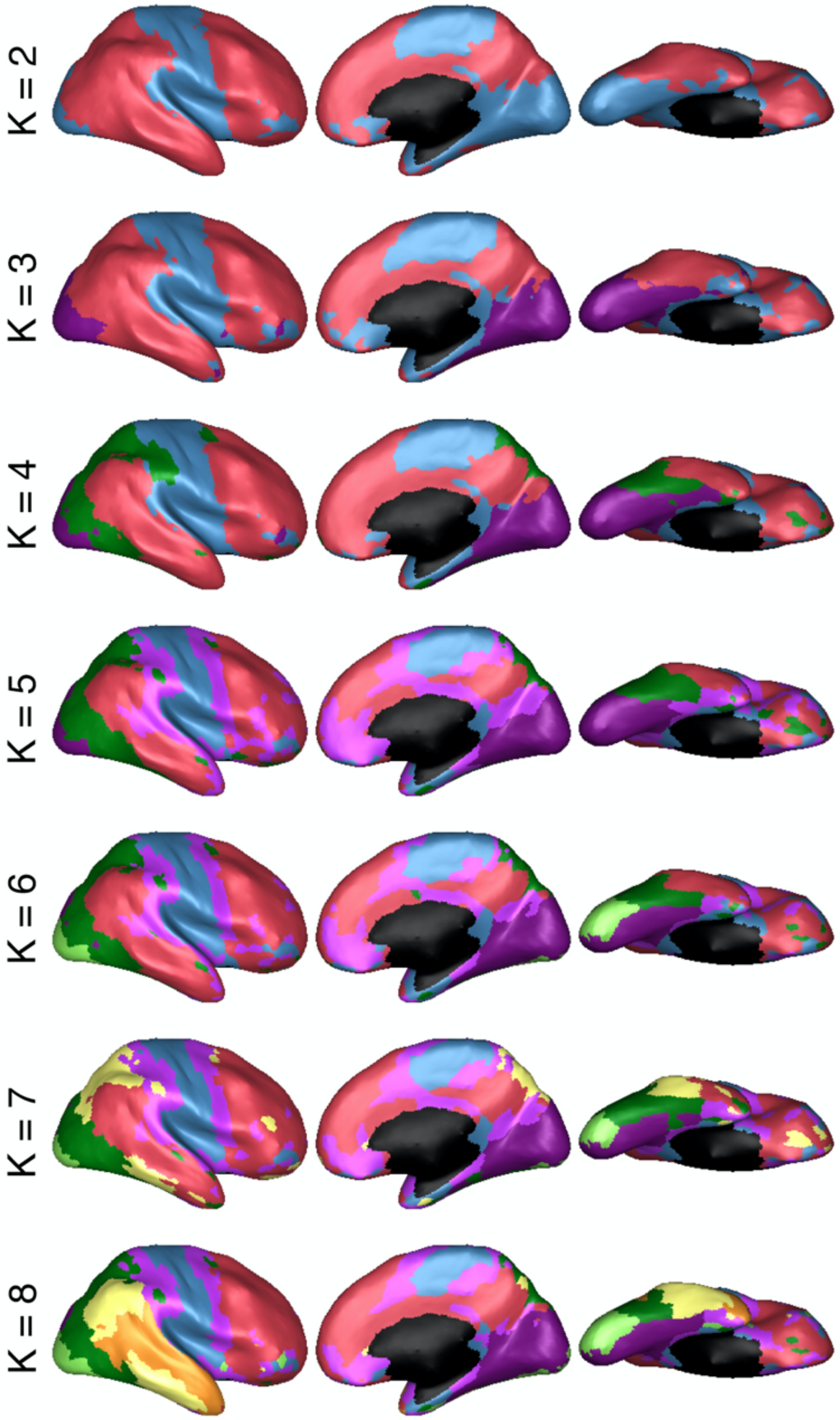


**Supplementary Figure 12**. Each row shows the cortex-only clustering solutions (on the test set of subjects) for an increasing number of clusters (k). Voxels are assigned a k-value based only on the similarity of their connectivity profile to all other voxels. All solutions are displayed on the right hemisphere for ease of visualization.

Higher clustering solutions further split the posterior networks and additional distinctions emerge in the ventral aspect. Note that only the cortex solutions show k solutions greater than 5, as those networks were unstable in the whole-brain solutions (not closely replicated in the test/train solutions). In the 6-network solution, a lateral component of the occipital lobe (light green) splits the visual network into two components. Then, in the 7-network solution, a temporoparietal network (yellow) emerges, though the boundaries of some networks shift in this less stable solution. Finally, in the higher-resolution optimal solution (the 8-network solution), the temporoparietal network components subsume the posterior default components seen in lower resolution solutions. Additionally, an orange network emerges, that is highly localized to the temporal lobe. In the main text, we discuss how these further subdivisions, particularly the gradient on the ventral surface of the brain may relate to higher-order visual function.

**Symmetry for Cortex-only Solutions by Network**

As stated in the main text, solutions were overall symmetric, with an average of 80% voxels with the same network assignment in the corresponding voxel across hemispheres. In both resolutions, the ventral attention network was the least symmetric (**Supplementary Figure 13**). Breaking up by network in the 5-cluster solution, most networks are comparable, but the ventral attention network is the least symmetric (mean = 0.66 for test and 0.64 for train). In the 8-cluster solution, the ventral attention network had the lowest symmetry (mean = 0.63 for test and 0.64 for train).

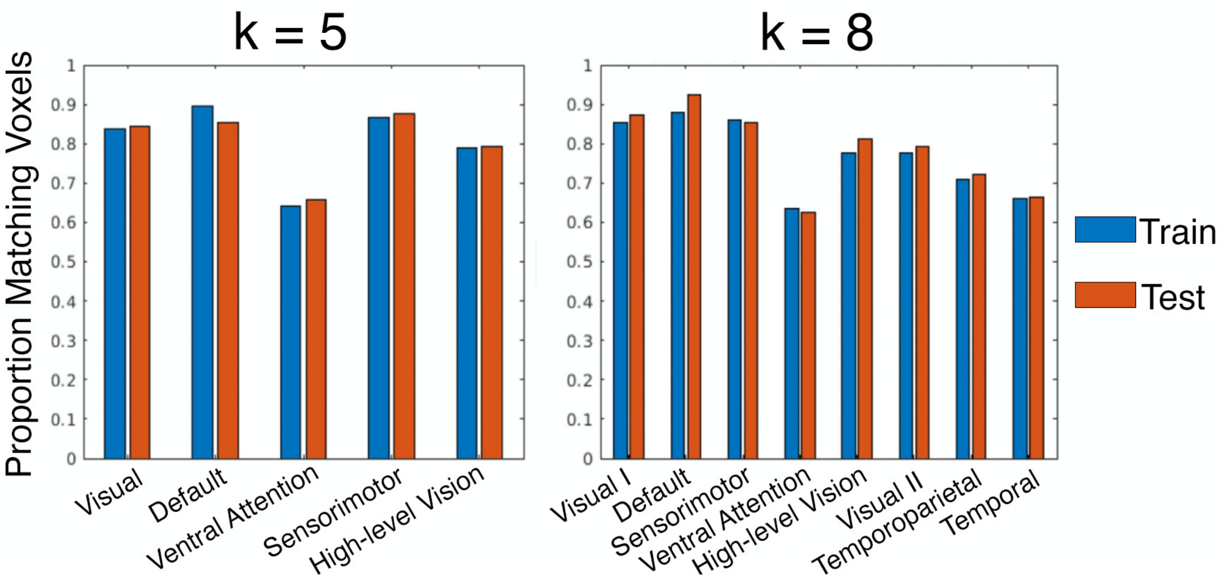


**Supplementary Figure 13.** Bar plots showing the symmetry by network in the test and train solutions for the cortical k= 5 (left) and k = 8 (right) solutions. Symmetry is quantified by the proportion of matching voxels which is the proportion of right hemisphere voxels that are classified as the same network as the network of the closest left hemispheric voxel.

**Individual Differences in All Optimal Solutions**

Figure 4a in the main text displayed the between-subject proportion of overlap for the cortical 5-network solution projected onto the surface. **Supplementary Figure 14** shows the same between-subject measure in the volume space for all three optimal solutions. In all three solutions, the highest between-subject agreement was in voxels in the sensorimotor network. Moderate overlap is seen in the occipital networks in all solutions as well.

**Comparison of 5-network Solution to Yeo et al., (2011)**

**Supplemental Figure 15** shows the quantitative comparison of the optimal 5-network neonate solution and the 7-network adult solution. The highest neonate and adult overlap were in the sensorimotor (dice = 0.74) and visual networks (dice = 0.69). The ventral and high-level neonatal networks had moderate overlap with the adult ventral (dice = 0.33) and dorsal (dice = 0.48) networks respectively. There were no neonate matches for the limbic or control network (dices = 0).

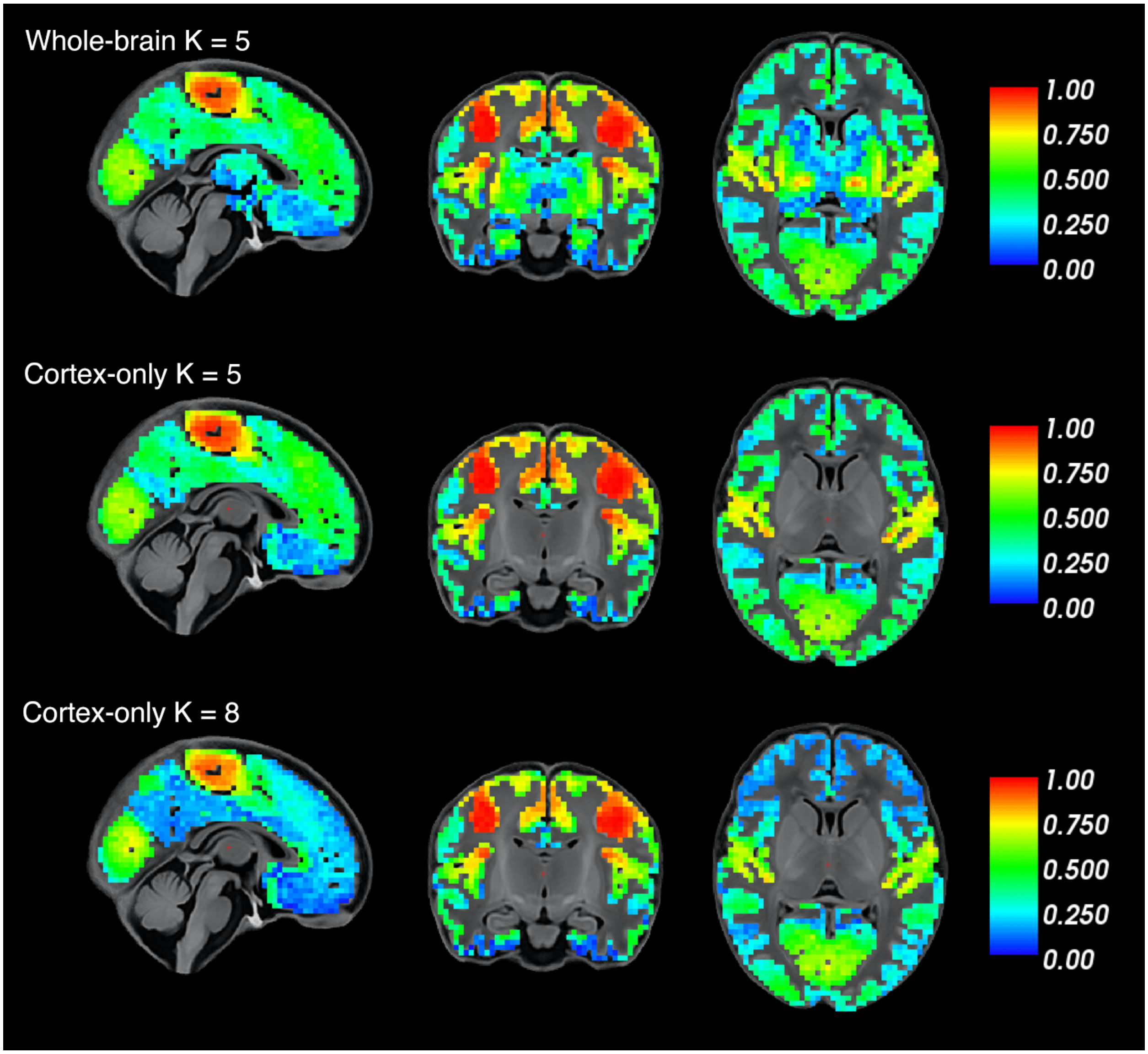


**Supplementary Figure 14.** Voxel-wise between-subject proportion of overlap for the three optimal solutions (rows). Voxels in warmer colors depict a higher proportion of individuals who were assigned to the same network.

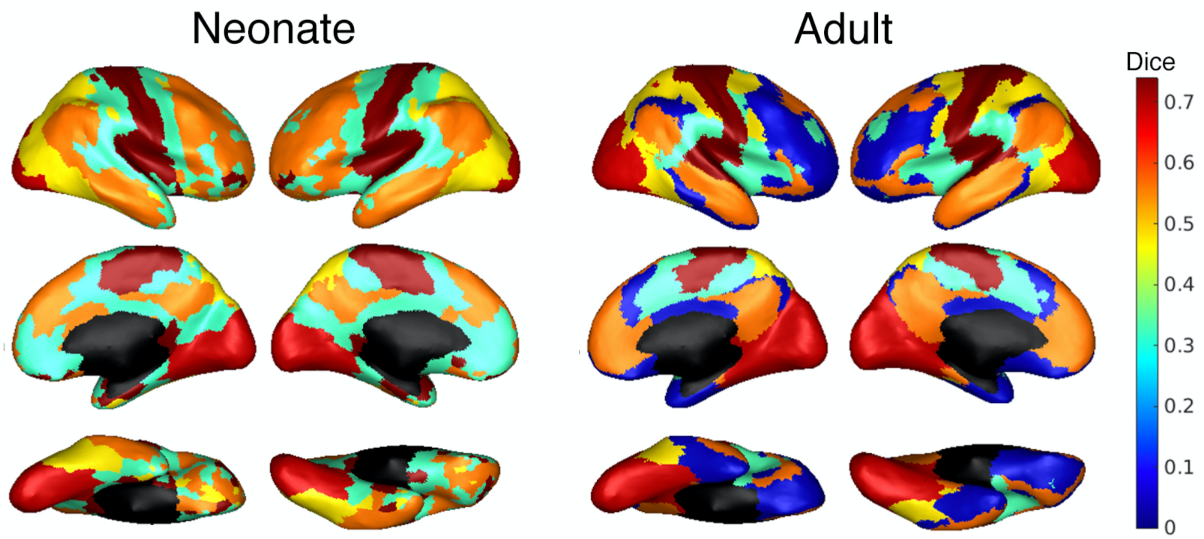


**Supplemental Figure 15**. Dice overlap between matched pairs of neonate (k = 7) and adult (k = 7) networks. Warmer colors show that a network is more similar between neonates and adults. The results are plotted onto the neonate networks (left) and the adult networks (right).

**Samples Matched to Networks**

The samples from the Allen Human Brain Atlas (AHBA) were matched to MNI coordinates for the ICBM 2009b nonlinear symmetric reference brain (Fonov et al., 2009). To get the samples in the FreeSurfer CVS average-35 MNI152 space, we aligned the volumes using FreeSurfer's mri_robust_register tool (Reuter et al., 2010). Each individual's samples MNI coordinates were then transformed into FreeSurfer space. We ensured all samples in each individual were successfully registered and were dilated before transformation if necessary.

These samples were matched to the cortex-only 5-network solution in MNI152 space. First, only samples in cortical gray matter were included, based on a cortical gray tissue mask from the FreeSurfer Desikan-Killiany parcellation (Desikan et al., 2006). For the left-hemispheric donors, samples were assigned to the subregion post-watershed transformation (see main text) if the sample was within that region. For the bihemispheric donors, in order to maximize the total number of samples included, cortical gray matter samples were matched to the closest subregion. **Supplementary Table 2** shows for each donor, the total number of samples released by AHBA, the number of samples matched to networks, and the total number of neonatal network post-watershed subregions with at least one matching sample. Because we average within an individual multiple samples within a single subregion, the total number of matched subregions also equals the total number of observations for that individual in the limma differential expression linear model.

**Supplementary Table 2**
***Tissue Samples per Allen Human Brain Atlas Donor***

| Donor ID | Hemisphere(s) Included | Total Samples | Matched Samples | Matched Subregions |
|---|---|---|---|---|
| H0351.1009 | Left | 363 | 92 | 73 |
| H0351.1012 | Left | 529 | 128 | 111 |
| H0351.1015 | Left | 470 | 119 | 102 |
| H0351.1016 | Left | 501 | 116 | 92 |
| H0351.2001 | Left and right | 946 | 675 | 254 |
| H0351.2002 | Left and right | 893 | 534 | 210 |

**Additional Enrichment Summaries**

Highlights of the functional enrichment analyses for the ventral attention, visual, default, and high-level vision networks are summarized in **Supplementary Tables 3—6**. Full enrichment results can be found in the **Supplementary Data**.

**Supplementary Table 3**
***Ventral Attention Enrichment***

| | p | q | Genes |
|---|---|---|---|
| Biological process | | | |
| regulation of nervous system development | $6.95x10^{-5}$ | $7.40x10^{-2}$ | HGF, DTNBP1, CPNE6, GPC4, ALK, PLXNA1 |
| neurotransmitter receptor activity | $3.51x10^{-7}$ | $6.97x10^{-4}$ | HRH1, HTR1A, HTR3A, HTR7, CNIH2 |
| neuron development | $6.43x10^{-4}$ | $2.95x10^{-1}$ | DOK6, HGF, PDGFB, DTNBP1, ACSL4 |
| neuron projection development | $8.64x10^{-4}$ | $3.49x10^{-1}$ | DOK6, HGF, DTNBP1, ACSL4, CPNE6 |
| regulation of neuron differentiation | $9.19x10^{-4}$ | $3.62x10^{-1}$ | HGF, DTNBP1, CPNE6, ALK, PLXNA1 |
| Cellular Component | | | |
| presynapse | $9.37x10^{-7}$ | $8.71x10^{-4}$ | DTNBP1, RAB3C, GPC4, CNR1, DOC2B |
| synapse | $2.94x10^{-6}$ | $2.19x10^{-3}$ | ACTC1, DTNBP1, RAB3C, GPC4, CNR1 |
| neuron projection | $4.49x10^{-6}$ | $2.78x10^{-3}$ | PCSK5, PDGFB, PDGFRA, CETN2, DTNBP1 |
| GABA-A receptor complex | $4.89x10^{-3}$ | $3.37x10^{-1}$ | GABRA3, GABRA5, GABRB1 |
| Disease | | | |
| Unipolar Depression | $4.59x10^{-6}$ | $3.02x10^{-2}$ | MAOB, TLR7, DTNBP1, ADCY7, ACSL4 |
| Depressive disorder | $5.02x10^{-6}$ | $3.02x10^{-2}$ | MAOB, DTNBP1, ADCY7, ACSL4, ALK |
| Generalized seizures | $1.34x10^{-5}$ | $3.02x10^{-2}$ | ADRA1B, CNR1, ALPL, HTR1A, PRKCD |
| Pain | $2.29x10^{-4}$ | $2.97x10^{-1}$ | HGF, HRH1, CNR1, HTR1A, HTR7 |
| Coexpression | | | |
| Genes with high-CpG-density promoters (HCP) bearing histone H3 dimethylation at K4 (H3K4me2) and trimethylation at K27 (H3K27me3) in brain. | $1.06x10^{-11}$ | $9.28x10^{-7}$ | DDAH2, PDGFRA, SPC25, IKZF1, WNT4, ADCY7 |

409 genes were found to be differentially expressed for the ventral attention network (Bonferroni corrected p-value < 0.01, log2 fold change < 0). We report the uncorrected p-value, the false detection rate (FDR) adjusted for multiple comparisons (q-value), and a subset of the first 6 overrepresented genes.

**Supplementary Table 4**
***Visual Enrichment***

| | p | q | Genes |
|---|---|---|---|
| Biological process | | | |
| Intracellular receptor signaling pathway | $2.61 \times 10^{-4}$ | $2.48 \times 10^{-1}$ | NCOA3, NRIP1, KDM5D, ALDH1A3, CALCOCO1, ARNT |
| Chromosome organization | $1.07 \times 10^{-13}$ | $8.99 \times 10^{-9}$ | NCOA3, KDM5C, SMC1A, JDP2, MCM2, KDM5D |
| Cellular Component | | | |
| Nuclear chromatin | $1.81 \times 10^{-21}$ | $1.50 \times 10^{-17}$ | NCOA3, NRIP1, KDM5C, JDP2, KDM5D, TRRAP |
| Nuclear body | $6.87 \times 10^{-15}$ | $2.86 \times 10^{-11}$ | NRIP1, HIPK1, AGO4, DUSP11, MPP1, USP36 |
| Disease | | | |
| Intellectual disability | $2.78 \times 10^{-8}$ | $3.48 \times 10^{-3}$ | HS6ST2, KDM5C, SMC1A, TRRAP, BRWD3, MEIS2 |
| Neurodevelopmental disorders | $8.60 \times 10^{-6}$ | $1.79 \times 10^{-1}$ | SMC1A, PPM1D, SETD2, CHD2, PHF2, PPP2R5D |
| Coexpression | | | |
| Human Messina_04_1962genes_TranscriptionFactors | $2.10 \times 10^{-28}$ | $2.30 \times 10^{-23}$ | NCOA3, NRIP1, KDM5C, JDP2, KDM5D, MEF2A |

2,243 genes were found to be differentially expressed for the visual network (Bonferroni corrected p-value < 0.01, log2 fold change < 0). We report the uncorrected p-value, the false detection rate (FDR) adjusted for multiple comparisons (q-value), and a subset of the first 6 overrepresented genes.

**Supplementary Table 5**
***Default Enrichment***

| | p | q | Genes |
|---|---|---|---|
| Biological process | | | |
| neuron development | $9.45x10^{-6}$ | $1.61x10^{-2}$ | USP9X, GPRIN1, SHC3, ADCYAP1, MRTFB, MIF |
| neuron projection development | $1.27x10^{-5}$ | $2.01x10^{-2}$ | USP9X, GPRIN1, SHC3, ADCYAP1, MRTFB, MIF, |
| generation of neurons | $1.61x10^{-5}$ | $2.30x10^{-2}$ | MAN2A1, USP9X, GPRIN1, SHC3, ADCYAP1, FZD8 |
| regulation of nervous system development | $2.91x10^{-4}$ | $1.60x10^{-1}$ | MAN2A1, ADCYAP1, MIF, RASAL1, ENC1, DBNL |
| Cellular Component | | | |
| neuron projection | $1.26x10^{-17}$ | $9.63x10^{-14}$ | MKKS, TTYH1, MORN4, USP9X, SYT9, ACTG1 |
| synapse | $6.99x10^{-15}$ | $2.67x10^{-11}$ | TTYH1, SYT9, ACTC1, ACTG1, ACTN2, ADCYAP1 |
| Disease | | | |
| Generalized seizures | $9.78x10^{-6}$ | $4.40x10^{-2}$ | ADCYAP1, ADRA1B, ALPL, NOS2, NPY, CCK |
| Anxiety Disorders | $5.81x10^{-6}$ | $4.40x10^{-2}$ | ACACA, MAOB, ADCYAP1, ADRB2, MIF, ALPL |
| Depressive disorder | $6.32x10^{-5}$ | $2.03x10^{-1}$ | MAOB, ADCY7, ADCYAP1, MIF, SLCO1C1, NOS2 |
| Coexpression | | | |
| Genes down-regulated in brain from patients with Alzheimer's disease. | $4.46x10^{-38}$ | $4.86x10^{-33}$ | MKKS, MAN1A1, USP9X, C3orf14, MDH2, TM6SF1 |
| Age down-regulated genes in the human frontal cortex. | $1.13x10^{-10}$ | $1.11x10^{-6}$ | ACTA1, MACIR, TAGLN3, ATP6V1A, CALM3, UBE2M |

287 genes were found to be differentially expressed for the default network (Bonferroni corrected p-value < 0.01, log2 fold change < 0). We report the uncorrected p-value, the false detection rate (FDR) adjusted for multiple comparisons (q-value), and a subset of the first 6 overrepresented genes.

**Supplementary Table 6**
***High-level Vision Enrichment***

| | p | q | Genes |
|---|---|---|---|
| Biological process | | | |
| N/A | | | |
| Cellular Component | | | |
| integral component of plasma membrane | $9.35x10^{-5}$ | $1.46x10^{-1}$ | KCNV1, MET, SLC22A18, ADRA1D, PCDH19, LRRTM4 |
| intrinsic component of plasma membrane | $1.67x10^{-4}$ | $1.46x10^{-1}$ | KCNV1, MET, SLC22A18, ADRA1D, PCDH19, LRRTM4 |
| Disease | | | |
| N/A | | | |
| Coexpression | | | |
| Genes with high-CpG-density promoters (HCP) bearing histone H3 dimethylation at K4 (H3K4me2) and trimethylation at K27 (H3K27me3) in brain. | $1.08x10^{-4}$ | $2.74x10^{-1}$ | CARTPT, NELL1, C1QL2, ADRA1D, NPTX2, COL12A1 |

85 genes were found to be differentially expressed for the high-level vision network (Bonferroni corrected p-value < 0.01, log2 fold change < 0). We report the uncorrected p-value, the false detection rate (FDR) adjusted for multiple comparisons (q-value), and a subset of the first 6 overrepresented genes.
